# Just Add Data: Automated Predictive Modeling and BioSignature Discovery

**DOI:** 10.1101/2020.05.04.075747

**Authors:** Ioannis Tsamardinos, Paulos Charonyktakis, Kleanthi Lakiotaki, Giorgos Borboudakis, Jean Claude Zenklusen, Hartmut Juhl, Ekaterini Chatzaki, Vincenzo Lagani

**Affiliations:** Gnosis Data Analysis PC, Heraklion, Greece; Department of Computer Science, University of Crete, Heraklion, Greece; National Cancer Institute, National Institutes of Health, Bethesda, MD, USA; Chief Executive Officer, Indivumed Group; Laboratory of Pharmacology, Medical School, Democritus University of Thrace; Institute of Chemical Biology, Ilia State University, Tbilisi, Georgia

## Abstract

Fully automated machine learning, statistical modelling, and artificial intelligence for predictive modeling is becoming a reality, giving rise to the field of Automated Machine Learning (AutoML). AutoML systems promise to democratize data analysis to non-experts, drastically increase productivity, improve replicability of the statistical analysis, facilitate the interpretation of results, and shield against common methodological analysis pitfalls. We present the basic ideas and principles of Just Add Data Bio (JADBIO), an AutoML technology applicable to the low-sample, high-dimensional omics data that arise in translational medicine and bioinformatics applications. In addition to predictive and diagnostic models ready for clinical use, JADBIO also returns the corresponding biosignatures, i.e., minimal-size subsets of biomarkers that are jointly predictive of the outcome of interest. A use-case on thymic epithelial tumors is presented, along with an extensive evaluation on 374 public biological datasets. Results show that long-standing challenges with overfitting and overestimation of complex non-linear machine learning pipelines on high-dimensional, low small sample data can be overcome.

## Introduction

The number of molecular biological **datasets** available (defined as collections of molecular profiles of several biological samples) is increasing at a rapid pace, presenting welcoming opportunities for new science. Public repositories such as Gene Expression Omnibus^1^, recount2^2^, Metabolomics Workbench^3^, and the NCI Genomics Data Commons^4^ collectively contain hundreds of thousands of datasets. The datasets are typically associated with an **outcome** (target, dependent variable) of interest, such as disease status, response to treatment, disease sub-type, quantitative phenotypic trait, and time to an event of interest (e.g., survival, complication, metastasis). Predictive models can be learnt (fit) to predict the outcome in future (out-of-sample) profiles, using modern statistical and machine learning methods. In addition, such methods can identify the (bio)**signatures**; in this context, biosignatures are defined as minimal-size subsets of molecular and other measured quantities that collectively lead to optimal predictions. Identifying the signatures, a task called **feature selection** (a.k.a., **variable selection** or attribute selection) in general, is a major tool for knowledge discovery, gaining intuition into the molecular mechanism of disease, identifying drug targets, or designing diagnostic assays with minimal measurements and cost. Despite the plethora of available data, algorithms, and computational power, a major bottleneck in molecular data analysis is still present: lack of human experts’ time. In addition, analyses are prone to methodological errors that invalidate results and mislead the community^5^. Is it possible to fully automate the sophisticated, advanced multivariate analysis of biological data? Can we democratize machine learning to life scientists and non-expert analysts? Can we reduce statistical methodological errors that creep into analyses?

As a response to the above challenges, the field of Automated Machine Learning (**AutoML**) recently emerged^6^. At a minimum, given the data and the outcome, an AutoML tool should return *a predictive model* and *an estimate of its predictive performance to out-of-sample predictions*. When it comes to translational medicine and molecular biology data the list of requirements and challenges increases. The first one is the ability **to statistically scale down to low sample sizes**. It is not uncommon for biological datasets to contain fewer than 100 samples: as of March 2020, 74.6% of the 4348 curated datasets provided by Gene Expression Omnibus count 20 or fewer samples. This is typical in rare cancers and diseases, in experimental therapy treatments, and whenever the measurement costs are high. A second challenge is the ability **to scale up to hundreds of thousands or even millions of features** (dimensions). Such high dimensional data are produced by modern biotechnologies for genomic, transcriptomic, metabolomic, proteomic, copy number variation, single nucleotide polymorphism (SNP) GWAS profiling, and multi-omics datasets that comprise of multiple modalities. The combination of a high number of features (*p)* and a low sample size *(n*), or as it is called “large *p*, small *n”* setting, is notoriously challenging as it has been repeatedly noted in statistics^7^ and bioinformatics^8,9^. A third challenge is to perform signature discovery, simultaneously with predictive modeling. The former is often the **primary** goal of an analysis: for example, for drug target discovery or precision pharmacotherapy, identifying the prognostic or predictive genes may be more important than the model itself. The latter serves only to quantify the predictive information contained in the signature. Reducing the number of features in a biosignature without compromising in sensitivity and specificity is a major demand in biomarker discovery and diagnostics.

In this work we describe the methodology of a web based AutoML platform to address the above requirements, called Just Add Data Bio or simply **JADBIO**, version 1.0.44 (March, 1, 2020) (Figure 1). The platform has been designed for non-experts to deliver high-quality predictive and diagnostic models employing standard, best-practices, and state-of-the-art statistical and machine learning methods. It identifies multiple (in cases of biological redundancy) equivalent signatures of predictive features. It scales up to hundreds of thousands of features, which implies that it can simultaneously consider multi-omics, clinical, and epidemiological features. JADBIO scales down to tiny sample sizes (e.g., 40) often encountered with molecular biology data, in the sense that it still manages to provide accurate, non-optimistic estimates of predictive performance.

**Figure 1.**
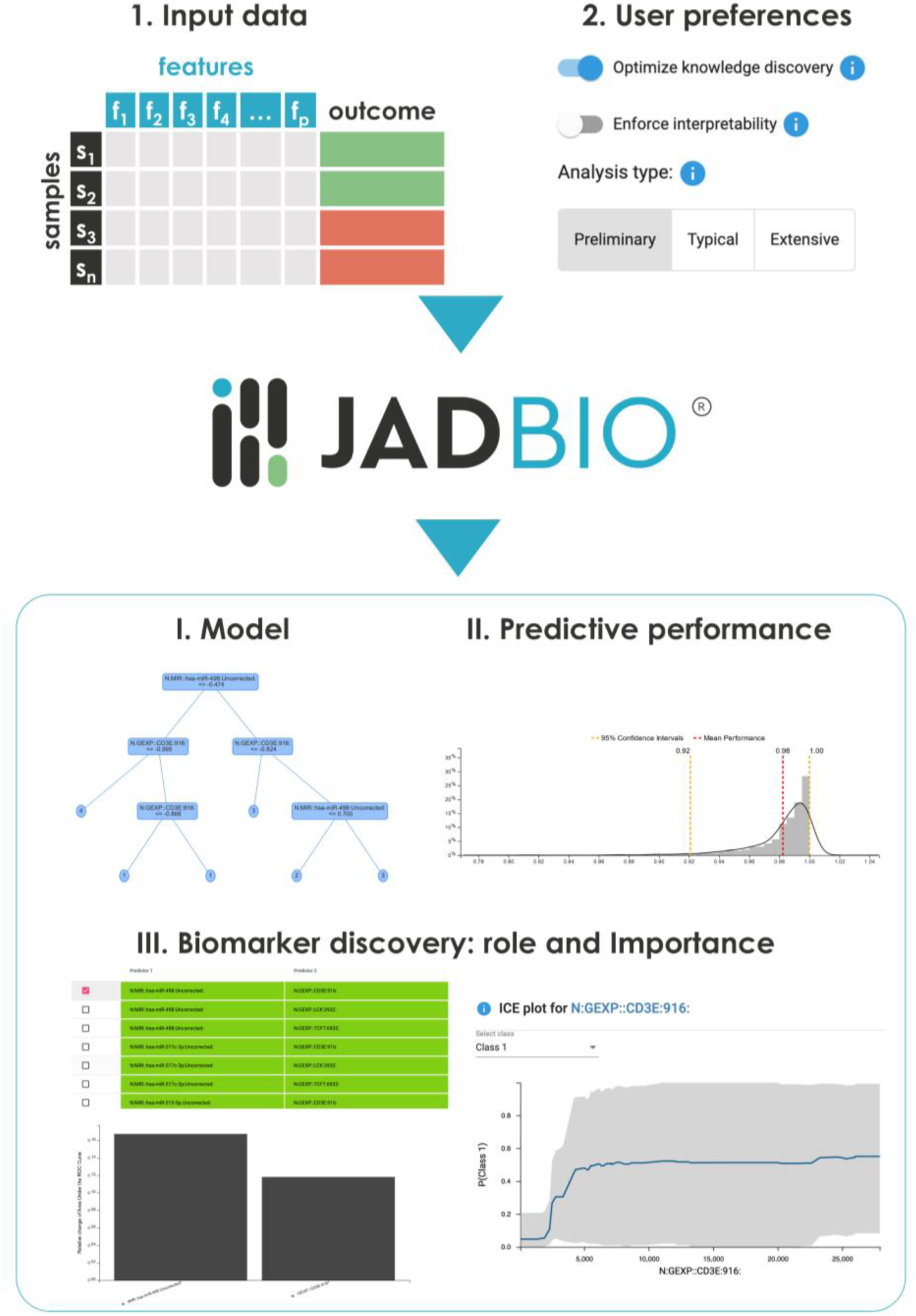
JAD Bio input-output scheme. JAD Bio takes as input a 2D matrix of data where the rows correspond to samples and columns to features (e.g., biomarkers). One of the columns is the designated outcome to learn to predict. It also accepts user-preferences. These can enforce or not biosignature discovery (optimize knowledge discovery), the interpretability of the final model, and define the time budget of the analysis (preliminary, typical, extensive). It then automatically produces a predictive model for a discrete (classification), continuous (regression), or time-to-event (survival analysis) outcome, estimates of its predictive performance, and multiple, equivalent, minimal-size signatures of biomarkers. Graphs and visual explain the impact and role of each marker in a signature.

JADBIO is specifically devised for life scientists without knowledge of coding, advanced statistics or data analysis expertise. It shields against typical methodological pitfalls in data analysis that lead to overfitting and overestimating performance and therefore to misleading results. The novel machine learning algorithms of JADBIO have been validated by the machine learning and statistical community^10–20^, while the system has produced novel results in nanomaterial property predictions^21^, suicide prediction^22^, speech classification^23^, bank failure prediction^24^, function protein prediction^25^, and breast cancer prognosis and drug response prediction^26^, to name a few.

In this paper, JADBIO’s user-centric functionalities are demonstrated on a use-case problem, discriminating between four Thymic Epithelial Tumors (TET, also-known-as thymoma) subtypes based on multi-omics data. In addition, the claims of correctness, as well as automation, are proven empirically on 374 public biological datasets, spanning 124 diseases and corresponding controls, from psoriasis to cancer, measuring metabolomics, transcriptomics (microarray and RNA-seq), and methylomics. The large-scale JADBIO evaluation answers several long-standing questions of the bioinformatics analysis community. It is shown that it is possible to avoid over-estimation on average in “large *p*, small *n”* settings, even when numerous, complex, non-linear machine learning models are tried. Robust (multi-variate) feature selection in a way that generalizes to new data is also possible; in fact, it is shown that in biological datasets there typically exist multiple, equally predictive feature sets. Finally, it is shown that a large percentage of public molecular datasets of all modalities does carry predictive yet untapped information about its corresponding outcome. Such information can lead to predictive models with potential clinical use by modern AutoML technologies.

## Results

### Use case on Thymoma Differential Diagnosis

A use case on thymoma tumors differential diagnosis is presented to demonstrate the functionalities, capabilities, and range of results and visuals of the proposed AutoML architecture. Results are sharable via automatically generated unique URLs. The ones for this analysis are found at https://app.jadbio.com/share/8292bad2-e130-4ef6-a5bc-b7cc727d5432. Primary tumor biopsies from one-hundred seventeen (117) thymoma patients were profiled by The Cancer Genome Atlas (TCGA, https://www.cancer.gov/tcga) for copy number variation, gene expression, methylation levels, microRNA and genomic mutations. Four (4) subclasses were defined in a previous study on the basis of all 27796 molecular measurements combined, using a multi-omics clustering approach^27^. The specific objectives of the use case are (a) to derive a classification model to disease subtype given as few as possible molecular features. Such a model can then be employed clinically to assess new thymoma patients with minimal measurement time, costs, and workload. (b) To validate the diagnostic accuracy of the model in terms of specificity/sensitivity on new subjects in order to evaluate its potential for effective clinical application. (c) To understand the role of the biomarkers in providing predictions.

The 2-dimensional 117×27797 data matrix with the measurements (rows corresponding to samples from individual patients, columns corresponding to features and the outcome) was uploaded to JADBIO (Figure 2a left). Subsequently, the user preferences and the outcome were specified (Figure 2a center) and the analysis began. In this example, the user preferences were set to “Extensive analysis” to extensively search for an optimal model at the expense of computational cost, and “optimize for knowledge discovery”, which implies enforcing feature selection (biosignature discovery). Once the analysis begins, JADBIO starts training models using different **configurations** on different sub-samples of the original dataset (cross-validation, see the Methods section). A configuration is defined as a machine learning pipeline that produces a predictive model given the data; it combines algorithms for transformations, imputation, feature selection, and modeling, along with their hyper-parameter values. The purpose of the search is to identify the configuration that produces the optimally predictive model and estimate its performance.

**Figure 2.**
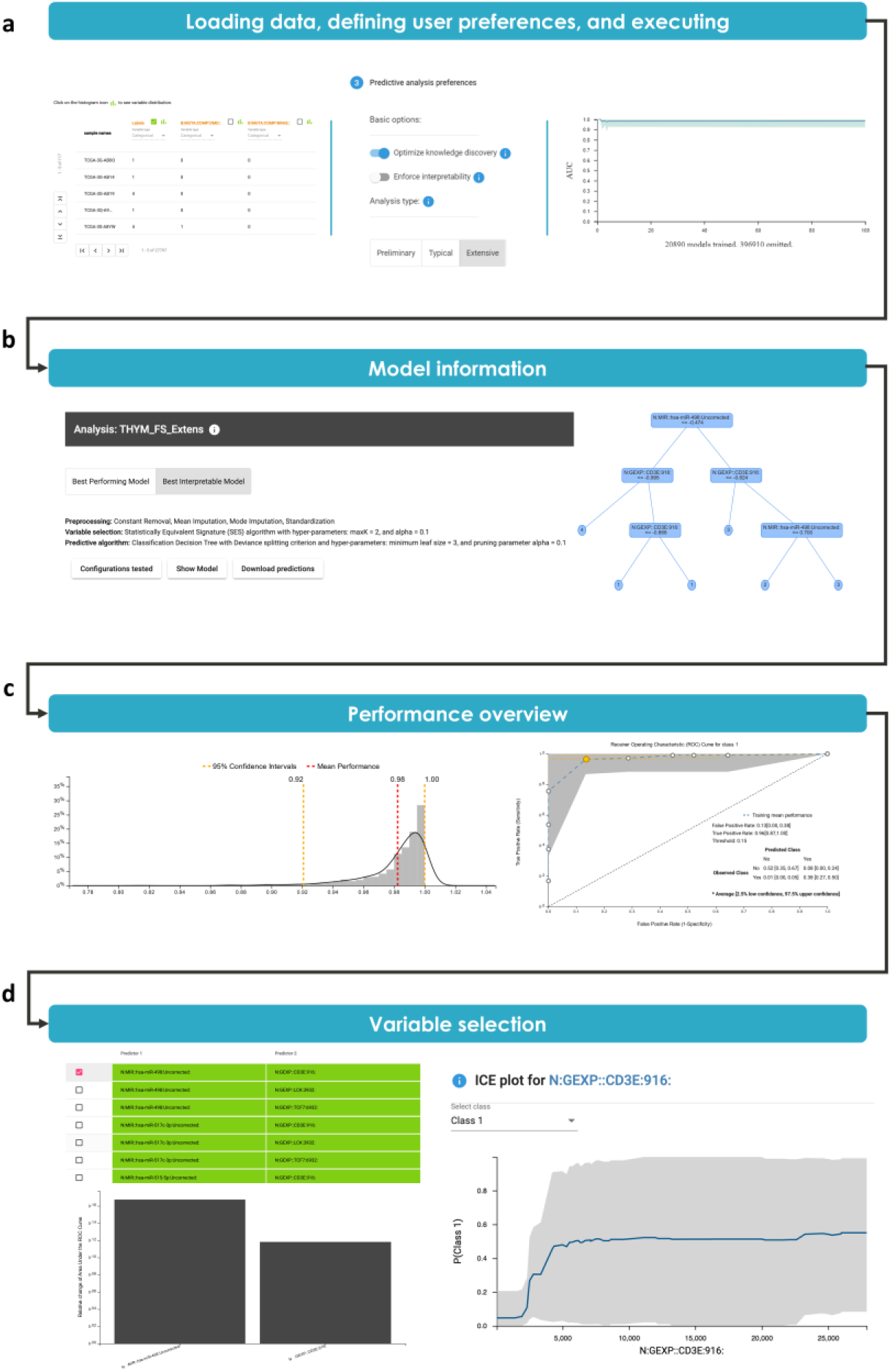
Thymoma Analysis – Performance and Selected Signatures. a) Loading the data (left) and selecting the analysis preferences (middle). The progress status of the analysis can be monitored in a progress bar plot (right). b) The winning configuration (pipeline) that produces the optimal, final model (left). The best model out of those that are humanly interpretable is also automatically produced (right). It is a decision tree that can classify to the four sub-types according to two selected features with a slight drop in predictive performance. c) The expected out-of-sample distribution of the AUC metric for the winning model (left). The ROC curve for the first subtype (class 1) against all the others (right). All metrics are adjusted for multiple tries and accompanied by 95% confidence intervals. The user can click on a circle (orange circle) to show the FPR and TPR and confusion matrix that corresponds to a model operating at that ROC point (see embedded table in the ROC curve). d) (top-left) Each row is an equivalent signature (equally jointly predictive set of biomarkers) for Thymoma sub-classes (7 out of 45 shown). Each signature comprises of two biomarkers. The importance weighting of each feature (left bottom). The Individual Conditional Expectation (ICE) plot of the CD3E biomarker; the plot indicates the average probability predicted for sub-type 1 given the value of the marker, expressed as RNA-seq reads (right). The predicted probability has a rapid jump around 2500 reads.

JADBIO’s AI system informs the user that it decided to fit at most 417800 models (not shown); by the end of the analysis, 396910 of these models were skipped as JADBIO determined that not sufficient progress was being made. During the analysis the user can peruse the progress made, the predictive performance of the best model found so far (bold blue line) and its confidence intervals (shaded areas, Figure 2a right). The final results depend on fitting 20890 models.

Once the analysis is complete the user can peruse the results. The steps that lead to the winning model are reported to the expert analyst (Figure 2b-left). In this case, the winning model is a Random Forest ensemble of 100 Decision Trees^28^ after feature selection with the SES algorithm^20^ is applied. JAB Bio also produces the winning model only out of those that are humanly interpretable, such as a single Decision Tree and generalized linear models (e.g., Ridge Logistic Regression). The practitioner can then gauge whether the drop in performance by using the interpretable model is significant enough to justify the use of a more complex model or not.

The Performance Overview tab shows the metrics of performance; for discrete outcomes (multiclass classification) these include sensitivity, specificity, precision, and recall, for each class individually, as well as the total accuracy and average (over all classes) area under the ROC curve (AUC). The distributions of each metric are also estimated (Figure 2c-left) from which 95% confidence intervals (CI) are derived. The CIs determine whether the model’s predictive performance is statistically significantly better than random guessing. Specifically, for the thymoma dataset, the AUC averaged over all classes for the winning model is 0.982 (C.I. 0.918 – 1.000) and balanced accuracy is 0.915 (C.I. 0.765 – 1.000). Both confidence intervals do not include the performance of random guessing (0.500 AUC and 0.250 balanced accuracy, respectively) and so are statistically significant. For comparison, the performance of the winning interpretable model, namely the Decision Tree (Figure 2b-right), is 0.968 (CI= [0.863, 1.000]), an arguably good enough approximation for most practical purposes.

**It is worth noting that all estimates are adjusted (controlled) for trying multiple algorithms (configurations)** and are conservative. This is true even for small sample sizes and high-dimensional data as in this use-case. This statement of correctness is proved empirically in the next set of computational experiments below. Without this adjustment, **the cross-validated accuracy of the winning model is overestimated**, as confirmed by our results on public data reported below. The statistical method that adjusts machine learning models’ performance for multiple tries and computes the confidence intervals is described in Tsamardinos et al. (2018)^12^ (a detailed discussion on this phenomenon is in the Online Methods).

Particularly for discrete outcomes, the ROC curves for each class against any other are presented (Figure 2c-right). The ROC curves show all achievable trade-offs between the False Positive Rate and True Positive Rate of the final model. Different trade-offs can be achieved by thresholding the probabilities for each class that are output by the model for clinical use of the model. For example, by classifying to Sub-type 1 any new sample predicted to be Sub-type 1 with probability higher than 0.15 (see embedded table in the ROC curve in Figure 2c-right) one can achieve a TPR (sensitivity) equal to 0.96 if they are willing to accept 0.13 false positives (FPR) on average. This trade-off and the corresponding confusion matrix correspond to the selected orange point on the ROC.

The *Variable Selection* tab summarizes the results of the variable (feature) selection, as shown in Figure 2d. The list of biosignatures found are presented (Figure 2d-top left). Each row corresponds to an equally well-performing signature (up to statistical equivalence). In this case, the four thymoma subclasses can be distinguished based on just two molecular features. The first such set of two biomarkers identified (called the *reference* signature) is the expression values of the gene CD3E and the miRNA miR498, respectively (see Supplementary Methods for markers’ names codebook). miR498 is the marker most associated (pairwise) to the outcome. However, CD3E ranks only 190 in terms of pairwise association with the outcome (p-values 10e-94 and 10e-56, respectively)! *In other words, if one performs standard differential expression analysis, they will have to select 189 other markers before reaching CD3E*. In contrast, JADBIO’s feature selection algorithms, recognize and filter out the redundant features. This example anecdotally illustrates that the most predictive signature is not always composed of the most differentially expressed quantities, but rather by predictors who complement each other in terms of informational content. It is actually possible *that markers with no pairwise association to (not differentially expressed by) the outcome to be necessary for optimal prediction*^29^. These examples illustrate the difference between standard differential expression analysis and signature discovery for predictive modeling.

For the CD3E gene, there exist 14 alternative choices of different gene expressions, while 2 other miRNAs can substitute for miR498. Each of the 45 corresponding combinations choices comprise a different but equally predictive biosignature (full list in Supplementary Table 1). Only the first seven combinations are shown in the figure. Figure 2d (bottom left) shows the individual importance for each of the two features of the reference signature (see Methods) which shows the drop in relative performance when a single feature is removed from the model. This is an illustration of JADBIO’s capability to identify multiple equivalent signatures. There are several reasons why multiplicity exists. It is not a contrived phenomenon but a commonly encountered one in biomedicine^30^. It is discussed in more detail in the context of the large-scale evaluation below.

In Figure 2d (right), the Individual Conditional Expectation (ICE) (see also Supplementary Methods and ^31^) for the CD3E is presented. The goal of the ICE plot is to explain the role of an individual feature in the prediction of the model. The y-axis corresponds to the predicted probability that a sample is of Sub-type 1 (i.e., P(Class 1)). The bold line in the plot shows how the average of the predicted probability as the observed values of the marker increase. The grey area indicates the variance of the predictions due to the values of the other biomarkers. In this case, the plot is like a step function: when the value of CD3E exceeds a certain threshold (around 2500 next-generation sequencing reads) the probability of a sample to belong to the first Sub-type 1 (Class 1) jumps higher. For low values of CD3E expression, the variance in the prediction is relatively smaller, indicating that when CD3E is low, the values of the other biomarkers do not affect the prediction as much. ICE plots facilitate interpretation of the role of a marker in the model, visualizing non-linear effects, and gauging the dependence of predictions on the combination of all features.

The *analysis visualization* panel of JADBIO presents different diagnostic plots according to the performed task. For example, Figure 3a shows a scatter plot of the profiles in the space of the two selected features in the first signature. It visually confirms that the four sub-types can be well-separated based on only these two features. It also makes obvious that non-linear models are required to classify the sub-types. The purpose such plots is to enable the user to visually inspect the results of the analysis and verify there are no obvious anomalies. For example, if in Figure 3a had a blue point (Class 1 sample) be placed well-within the green points region (Class 2) it could indicate a mis-labeled or corrupted sample. Once a satisfactory model is produced, one may decide to apply it in a clinical setting. The application of the model to an external validation set or on a new profile is also automated: Figure 3b shows the generated web-page (micro-service) that implements the thymoma model. The form accepts the values for the selected features of the new profile and outputs the estimated probabilities for each class.

**Figure 3.**
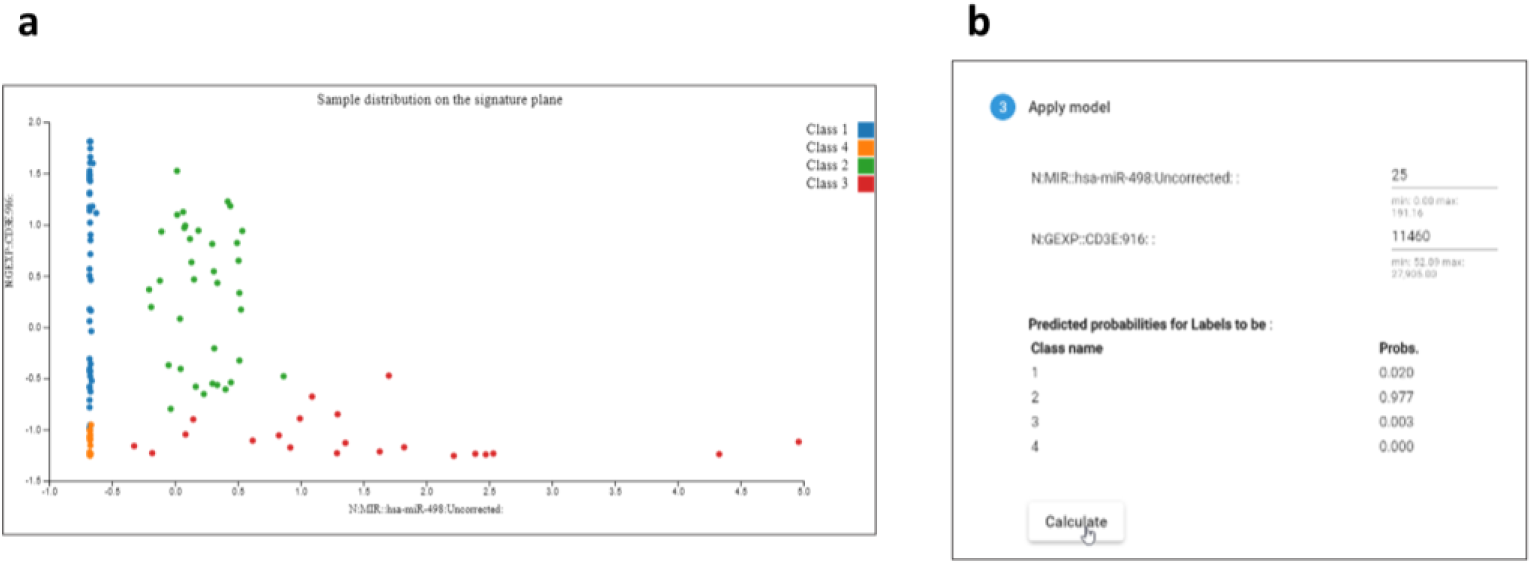
Thymoma Analysis – Model Inspection. a) Scatter plot of samples based on the two selected molecules (features) of the reference signature identified. The four thymoma subclasses are well-separated, visually confirming the performance metrics. Separating the four subtypes obviously requires non-linear methods. b) Applying the final model to an external validation set or on a new profile in a clinical setting is also automated through the JAD Bio graphical interface, as executable code, or through an API call.

The computational time required for the analysis of this dataset largely depends upon the user preference settings and the number of CPUs employed. When using three computational cores executing in parallel, the Extensive setting requires approximately eight hours for fitting 20890 models out of the 417800 models planned initially. This is because JADBIO employs heuristics to stop computations early if progress in terms of performances and accuracy of estimations has plateaued. Repeating the analysis setting the time budget to “Preliminary” the user gets very similar results for this problem (same best model and approximately the same performance estimate) within only eighteen minutes: in this task, the extra search for an optimal model does not pay out. Finally, we’d like to note that, while the thymoma problem is a multi-class problem, JADBIO can also handle continuous outcomes (e.g., a quantitative phenotypic trait, or expression level of a gene), as well as, time-to-event outcomes (e.g., time to death, metastasis, relapse, or complication). The functionality of JADBIO can also be called through an Application Program Interface (API) for batch processing and analyses of datasets (this functionality made possible to produce the extensive experimental results in the next section).

### Large-Scale Evaluation on Public Data

Computational experiments have been performed on a large corpus of 374 high-dimensional dataset, including transcriptomics, epigenomics, and metabolomics data. The datasets are related to 124 different diseases, including chronic obstructive pulmonary disease, psoriasis, mental health diseases, and with diseases of cellular proliferation (i.e., different types of cancers) being the most represented group with 157 datasets. In total, we analyzed 38901 samples (molecular profiles), corresponding to ~3×10^9^ data values, with each profile measuring between 6 and 485514 molecular quantities (variables, features), 82300 on average. For this computational experiment, 45,766,839 predictive models were constructed. Supplementary Table 2 shows the characteristics of the datasets involved in the experimentation. Curated datasets for binary classification were obtained from online repositories, such as BioDataome^32^ (transcriptomics, methylation data) and Metabolomics Workbench^3^ (metabolomics). Dataset selection and preprocessing, as well as the experimental protocol, are described in detail in the Methods section. In short, each dataset is split in two subsets; JADBIO uses the first half for training and the second for testing (application of the model to out-of-sample data). The process is repeated by exchanging the role of the training and test sets for a total of 748 JADBIO runs. After each training, JADBIO returns the best predictive model along with an estimate of its performance (point estimate and confidence intervals). The accuracy of the estimate is then assessed on the other half of the data (holdout set). We use standard bootstrapping to produce the confidence intervals of the hold-out predictive performance.

### JADBIO avoids overestimating performance

For translational applications of machine learning and statistical modeling it is important not to overestimate the expected predictive performance of the models. This is a particularly challenging task given the low sample size, the high dimensionality of the problems, and the fact that modeling is combined with feature selection. In addition, *when trying to optimize and tune the algorithms one must adjust performance of the winning model for multiple tries*. JADBIO’s estimation protocol, however, addresses all these challenges. In the upper panels of Figure 4, we plot JADBIO’s out-of-sample AUC estimates obtained from the training data with the bootstrap corrected cross-validation (BBC-CV) algorithm^12^, versus the actual AUC achieved by the application of the model on the hold-out test data (blue points). Each point corresponds to a dataset and an estimation. Points *above the diagonal underestimate* performance (training estimate is smaller than the actual on the hold out) and *points below the diagonal overestimate* performance. The blue bold line shows the average behavior obtained using a locally estimated scatterplot smoothing (LOESS), see Cleveland et al. (2017)^33^. The lower panels present the same results grouped according to different levels of predicted performance estimated on the training set. For each group the average bias (actual minus predicted AUC) is reported along with its corresponding standard deviation (error bars).

**Figure 4.**
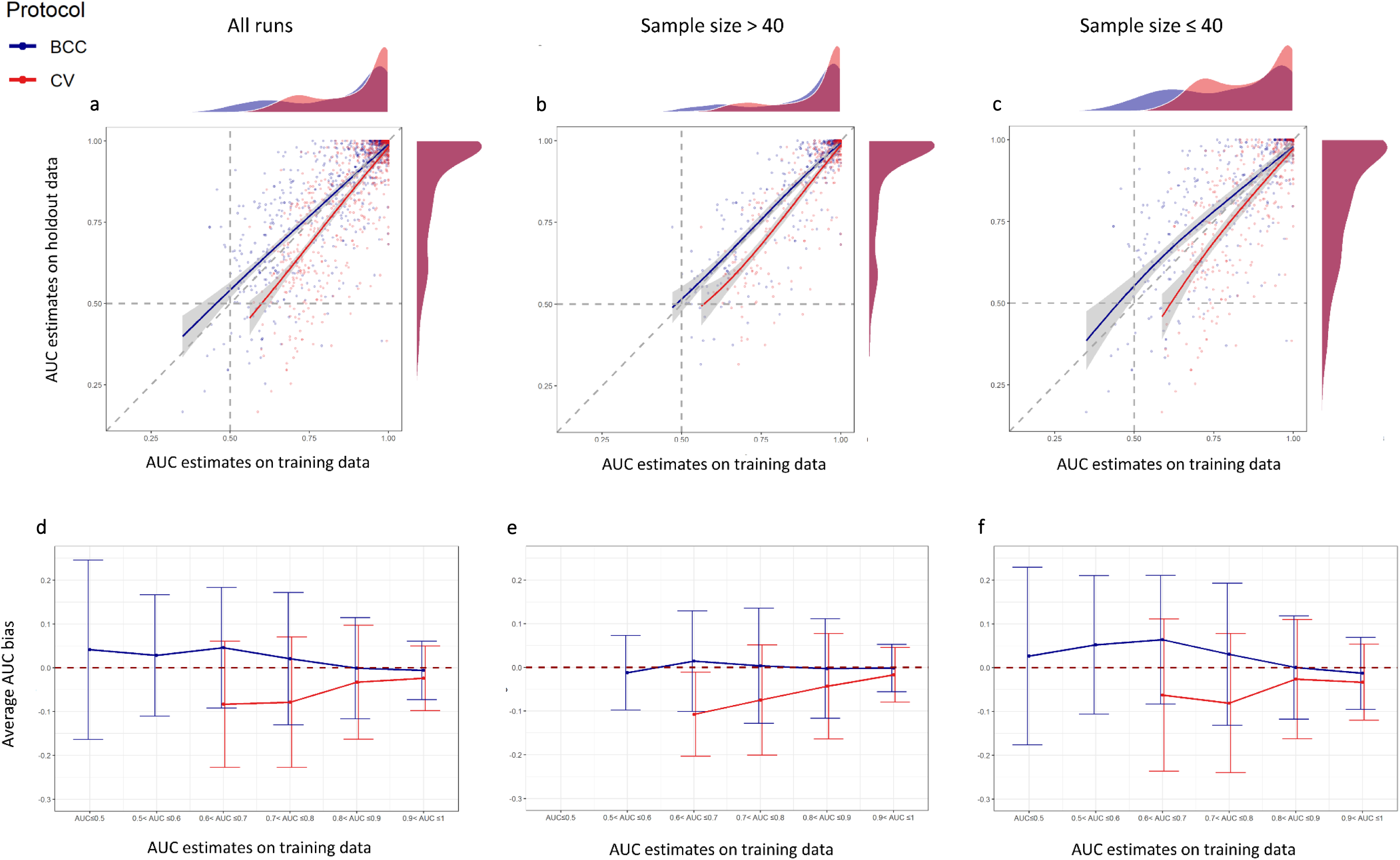
JADBIO predictive performances on classification tasks. The upper panels contrast the Area Under the receiver operating characteristic Curve (AUC) estimated on the training set (x-axis) against the AUC independently computed on the holdout set (y-axis). The leftmost panels include all JADBIO runs, while the center and rightmost panels respectively display runs where the sample size of the training set is above or below the median value (40 samples). Each run is represented by two dots: a blue one where the training cross-validated AUC estimate is conservatively adjusted for multiple tries with a bootstrap-based algorithm, and a red one where it is not. Loess regression on each set of dots is represented with a line of the same color (95% confidence interval reported as grey area). Markers over the diagonal dashed line indicates conservative estimates, while the area below the diagonal line contains overoptimistic estimates. JADBIO produces reliable estimates on average (blue line on or above the diagonal). Unadjusted cross-validation estimates overestimate performance (red line below the diagonal). The lower panels summarize the same information by binning runs according to performance levels on the training set (x-axis). Bins containing less than five runs were excluded. The y-axis reports the average bias (actual minus estimated performance), with error bars denoting the ± standard deviation intervals. The dark-red dashed line at zero indicates the absence of bias; points over this line marking an average conservative behavior and points below the line denoting over-optimist performance estimates. This representation confirms as well that JADBIO estimates are more reliable on average than the ones produced by uncorrected crossvalidation. While the variance of both protocols is non-neglectable, the uncertainty is reduced for bins corresponding to higher expected performance.

Across all datasets (Figure 4a), the Spearman’s rank correlation between training estimates and hold out performance is 0.815 (*p*-value < 2.2e-16). The average bias (actual minus estimated performance) is 0.008 AUC points, meaning that *performances are slightly underestimated*. We also note that even though the average behavior is close to ideal, for each individual dataset there is significant variance in the estimation, as some points fall far from the diagonal line. The variance is higher for the smallest sample sizes (≤ 40, stand. dev. 0.128) and smaller for larger sample sizes (> 40, stand. dev. 0.081), as shown in panels b and c of Figure 4, respectively. Based on the results, *we argue that accurate, non-optimistic estimations of advanced machine learning models are possible even in small-sample, high-dimensional settings*.

Referring again to Figure 4, the red dots denote estimations produced by cross-validation but without the BBC adjustment (standard cross-validation, CV). The average behavior (shown in the red, bold line) clearly shows that *on average, (unadjusted) cross-validation performances of the winning model are overestimated*. These findings on biological data corroborate the ones in Tsamardinos et al. (2018)^12^ on general types of data. The mean bias is −0.04 (stand. dev. 0.109), indicating that on average the training-set estimate is four points higher than the actual AUC value. Small-sample studies (≤ 40) again exhibit an increased bias and a higher variance in estimation, equaling −0.047 and 0.128 respectively. While −0.04 AUC overestimation on average may sound acceptable, panels D, E, and F present a more detailed and clearer picture: *the lower the estimated performance, the higher the overestimation and variance of the simple, unadjusted CV*. This is not an accident as the AUC can be considered the success probability of a binomial distribution, which has the highest variance of estimation at 0.50. Specifically let us focus on the bias when AUC is estimated to be within [0.6, 0.7] and [0.7, 0.8] in panels D, E, and F: it ranges around −0.1 and can also easily reach −0.2. Overestimations are even worse for lower sample sizes (panel F). *This translates to a practitioner estimating the AUC of the winning model using the standard (unadjusted) CV as 0.7, when in fact it equals random guessing (0.5 AUC)*. In summary, adjusting for the bias of CV estimate of performance for multiple tries is necessary for small sample sizes and low estimate values. On the other hand, high estimates of classification performance are typically more reliable.

In more than half (61.5%) of JADBIO runs, the AUC estimates in both the training and the test sets were above 0.80, indicating substantially informative results. In 66.7% of the runs the lower bound for the AUC confidence interval is above 0.5 for both training and holdout set, meaning that both estimates are better than random guessing in a statistically significant way. No significant deviation from this general trend is observed when the results are broken down according to different omics types. The lowest performances are registered for metabolomics data, where still 62.0% of the runs present statistically significant AUC in both the training and the holdout set, while RNA-seq seems to be the most informative technology in the sense that most analyses return models with AUCs being statistically significantly different than random guessing in 74.1% of the times.

*The results support that in general, omics datasets of different types do contain useful diagnostic and predictive information*; *this information can be extracted in the form of potentially non-linear predictive models using AutoML architectures even when sample size is less than 40 samples*.

### JADBIO accurately estimates statistical significance

AUC estimates that are statistically significant at the 0.05 level on the training set (i.e., with 95% confidence intervals above 0.50 AUC) are followed by statistically significant AUC values on the holdout set in 96% of the cases. This means that the confidence intervals provided by JADBIO allow one to successfully identify models that are likely to perform better than random guessing on new data.

### JADBIO selects parsimonious yet highly predictive biosignatures

We next evaluate the impact of feature selection on JADBIO predictive performances. Specifically, we contrasted the JADBIO results obtained by enforcing feature selection versus the results obtained by using all features as predictors (Figure 5a). We observe that models based on minimal-size signatures are only slightly less predictive than the ones based on all features: the median difference in holdout AUC is 0.023. At the same time, the compression ratio achieved by JADBIO (defined as the number of selected predictors over the number of initial variables) peaks around four orders of magnitude, indicating that only a handful of predictors are usually selected out of tens of thousands (89.3% of the datasets had signatures containing 5 or fewer biomarkers). *These results support the hypothesis that the predictive and diagnostic information of high-dimensional datasets for typical outcomes can be summarized by just a few biomarkers*.

**Figure 5.**
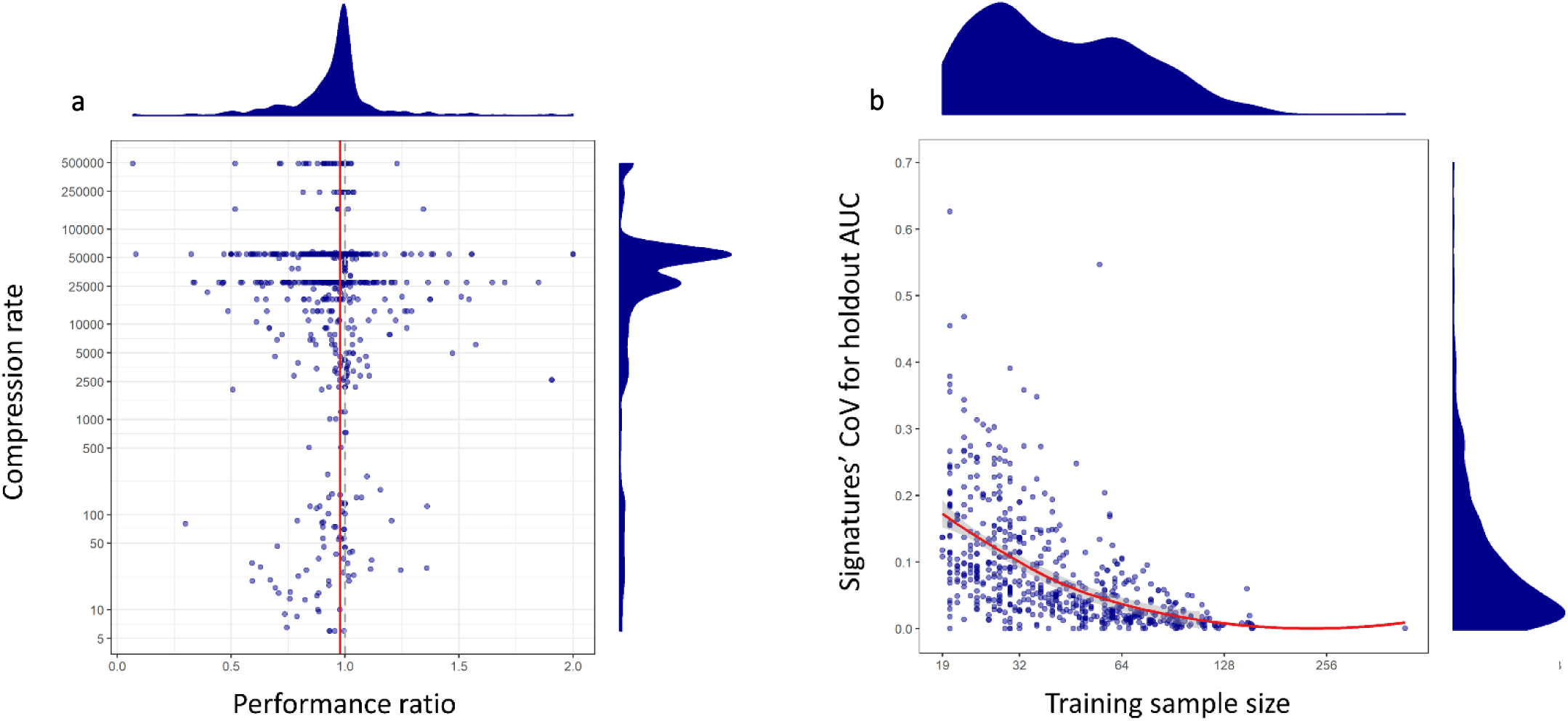
JADBIO knowledge discovery capabilities in classification tasks. A) Predictive performances with and without feature selection. The x-axis reports the ratio between the holdout AUC value obtained using feature selection versus the value obtained not using it; values above one indicate feature selection outperforming the use of all candidate predictors. The y-axis reports the level of compression, computed as the ratio between the number of selected features versus the number of all candidate features. The higher a point on the y-axis, the larger the compression obtained in the corresponding JADBIO run. The dashed vertical line indicates equal performances for the two approaches (ratio = 1), while the red solid line indicates the actual median ratio value (0.977). B) Relation between the number of samples in the training set (x-axis) and the coefficient of variation (CoV, standard deviation over mean) for the holdout AUC computed across equivalent signatures. Each point represents a JADBIO run where multiple signatures were identified; feature selection was enforced in all analyses. The trend followed by the data is estimated with Loess regression (red solid line; 95% confidence interval as grey area).

### Multiple, equivalent signatures are frequent in molecular biological data

When feature selection is enforced, JADBIO identifies more than one signature in 628 (84%) runs, while more than 10 signatures are retrieved in 440 (59%) of the runs. The prevalence of the presence of multiple equivalent signatures in biological data has been noted before^30^ and is corroborated by the present study. There are several reasons explaining multiplicity. First, when sample size is finite, the true best signature cannot be statistically distinguished from other candidates. Second, biological systems have built in redundancy which leads to statistical informational redundancy and multiple statistical signatures.

The number of identified signatures does not seem to depend upon the size of the training set, at least for small to medium sample sizes (Supplementary Figure 1). To assess whether the signatures indeed obtain statistically equivalent predictive performances, for each run we computed the Coefficient of Variation (CoV), defined as the ratio between the standard deviation and the average of the AUC values from equivalent signatures. Out of the runs with more than a single signature, roughly half (298, 47%) have a CoV below 5%, and 450 (71.7%) below 10%, indicating relatively narrow distributions for the AUC values estimated by equivalent signatures (Figure 5b, density plot for the y-axis). The CoV for AUC values from equivalent signatures is clearly inversely proportional to the logarithm of the sample size (Figure 5b). This indicates that determining actual equivalence between predictors is more challenging in small samples, most likely as a combined effect of (a) diminished statistical power in the training cohort and (b) higher variance in the performance estimation in the holdout set.

Overall, *the results support the claim that multiple optimal equivalent signatures are common in typical predictive tasks in molecular biological data*.

### Non-linear models are often optimal

Next, we examine whether non-linear models offer an additional predictive performance boost or not. While non-linear models can capture a wider set of types of associations and patterns in the data, they also tend to overfit. Therefore, linear models have been found to often perform better with lower sample sizes. This phenomenon is related to the bias-variance decomposition in pattern recognition and statistics^34^. However, in our experiments, Linear Ridge Regression was selected only 4.41% of the time as optimal, while linear Support Vector Machines (SVMs) were selected 6.82% for a total of 11.23% for linear models. Notice that the latter is a linear, but not a statistical based (parametric) model; it is a machine-learning type of model. Non-linear models were selected with the following frequencies: Decision Tree 4.41%, Random Forests (RF) 49.47%, Gaussian or polynomial kernel SVMs 34.89%. Notice that RFs and SVMs maybe nonlinear models however, they have strong mechanisms to shield against overfitting (bootstrapping and structural risk minimization, respectively). Our findings corroborate the ones reported in an extensive evaluation of 179 classifiers ^35^: RF, polynomial, and Gaussian kernel SVMs are often the top classifiers. When JADBIO was restricted to report only humanly interpretable models, Decision Trees were selected 24.73% of the times, while Linear Ridge Regression was selected 75.27% of the times (no other interpretable models were considered). Our results show that: *(a) modern non-linear models (RFs, SVMs) are often optimal for the analysis of low-sample, molecular data. (b) Optimizing the choice of the classifier is meaningful, as there is no single classifier that dominates all others in any dataset*.

### JADBIO achieves competitive predictive performance vs auto-sklearn

Auto-sklearn^36^ is among the state-of-art tools for automatic machine learning, winner of both international AutoML challenges competed so far^37^. In contrast to JADBIO, auto-sklearn is not specifically designed for the needs of biomedical researchers. It focuses on producing a predictive model, which is rarely the only piece of information required by a life scientist (see Xanthopolous et al.^38^ for a description a list of dozens of user-centric requirements and criteria of modern AutoML platforms other than predictive performance). Importantly, it does not even return a point estimate of the predictive performance of this model; the user is advised by its authors to use a separate hold-out set to estimate performance. These samples are “lost to estimation” which is unacceptable in many typical biomedical applications where the samples are scarce. Deciding on the size of the hold out also requires data analysis expertise, which implies that auto-sklearn cannot be used by non-experts. Auto sklearn does not return confidence intervals of performance which is important for clinical applications of the models: one cannot apply a model with 0.90 AUC (point estimate) in clinical practice if the variance of this estimate is within a confidence interval of [0.6, 0.95], for example. Feature selection is often *the primary task* in biomedical applications. JADBIO emphasizes feature selection and in addition, it performs multiple feature selection, i.e., identifies multiple feature subsets that are equally predictive. Presenting these feature subsets is quite important to biologists who try to understand the underlying biological mechanisms of the system under study. It is also important to biotechnologists who are designing diagnostic assays and would like to reduce the measuring cost. In contrast, auto sklearn does not perform feature selection, let alone multiple feature selection. In fact, its design inherently obfuscates feature selection: the selected model is an ensemble of 50 models (which in turn could be ensembles such as Random Forests) and each single model could be employing different subsets of features. On a conceptual level, we would categorize auto-sklearn as a *general purpose CASH library* (combined algorithm selection and hyperparameter optimization^39^), while JADBio is a AutoML *platform specializing to the analysis of molecular data*.

On a quantitative level, we attempted to repeat all classification analyses with auto-sklearn; however, the auto-sklearn system successfully finished only 453 (60.56%) runs out of the 748 completed by JADBIO. The remaining runs either stopped due to timeout (106 runs, 14.17%) or generated an internal exception (189 runs, 25.27%). Particularly, auto-sklearn managed to analyze only 14 runs out of 60 for the methylation data, which are characterized by the highest dimensionality (485514 measurements). This is somewhat expected, given that auto-sklearn is not specifically designed for low-sample, high-dimensional data. We next computed the ratio between the holdout AUC values produced by JADBIO and auto-sklearn on common successful runs. The average of the resulting distribution is statistically indistinguishable from 1 (average: 0.99, one-sample t-test p-value 0.347), indicating that the two systems perform equally well. JADBIO produces interpretable (≤ 25 elements) signatures on 23.52% of the runs while auto-sklearn models always employs the whole set of initial predictors. Requiring JADBIO to consider only the configurations including feature selection leads to AUC values that are on average 6% lower than the auto-sklearn results, however by using ≤ 10 variables in 96.47% of the times and ≤ 25 in all runs (Supplementary Figure 2). *These results indicate that JADBIO performs on par with autosklearn in terms of predictive performances, often offering results amenable for inspection and knowledge discovery*.

## Discussion

While Big sample Data present computational challenges, *small-sample data present statistical estimation challenges*: overfitting, overestimating, and model selection difficulty are exacerbated phenomena. Previous studies have already underlined these difficulties^40^, showing that not taking these issues into consideration can result in the publication of over-optimistic results^9^. In translational and biological research, additional challenges are present. A predictive model alone is hardly ever satisfying to practitioners. Interpretability and intuition into the molecular mechanisms involved in the prediction are required. This is often achieved by *signature identification (feature selection)*, i.e., minimal-size subsets of biomarkers that are jointly optimally predictive. It is a notoriously difficult problem. It is not only irrelevant markers that need to be filtered out but also redundant ones. In addition, markers that are not predictive in isolation (high unconditional p-value) may become predictive when considered in combination with other markers. To make things worse, prior research has shown that multiple such signatures are typically present in molecular data. Ideally, one would like to report *all signatures* that lead to optimal models (up to statistical equivalence) so as not to mislead the clinician or the biologist and provide choices to the designer of diagnostic assays. This is the *multiple feature selection* problem, as it is called, and it is only recently that its importance is gaining recognition^30^. Another challenge is the automation of the analysis, end-to-end. The algorithms to apply and choices of hyper-parameters depend on the size of data, the type of the features and the outcome, the sampling methodology (matched case-controls vs. cohort data), and the user preferences (favor interpretability over predictive performance). An expert analyst knows which algorithms may be promising and which are statistically inappropriate or computationally intractable in each case. An automated system needs to encode similar knowledge to make reasonable decisions in all cases.

In this work, we present a system for automated machine learning, called JADBIO, that addresses all the above challenges. An extensive evaluation on a compendium of 374 omics datasets spanning 124 diseases clearly shows that models created by JADBIO generalize to new samples as estimated by the training data. The confidence intervals reported accurately assess performances significantly better than random guessing in out-of-sample predictions. JADBIO identifies relatively small signatures of a few markers when enforcing feature selection (<5 markers out of a total of 82300 markers on average, in 89% of the datasets) by sacrificing 0.023 AUC points (median drop in performance) vs. models that may employ all available features. JADBIO discovers more than 10 multiple signatures in 59% of the runs, indicating signature multiplicity is a commonly occurring phenomenon. Non-linear models (Random Forests, non-linear SVMs, and Decision Trees) are selected as best in 88% of the time, suggesting that modern analysis of biological data should not be limited to standard, linear, statistical models. Finally, it is worth noting that in 61.5% of the analyses the AUC estimates on both the training and the test sets were above 0.80 across omics types. Molecular omics data do contain useful diagnostic and predictive information with high potential for the clinical practice. Their full exploitation has so far been impeded by lack of human expert time, as well as the statistical challenges to overcome.

What are the key ideas that enable JADBIO to overcome the challenges? The use of an AI decision-support system *that encodes statistical knowledge about how to perform an analysis* makes the system *adaptable* to several data sizes, data types, and user preferences. An automated search procedure in the space of appropriate combinations of algorithms and their hyper-parameter values, trying thousands of different machine learning pipelines (configurations) *automatically optimizes choices*. Protocols that estimate the out-of-sample predictive performance of each configuration, particularly the repeated, stratified, cross-validation protocol suitable for small sample sizes *reduce the uncertainty* (variance) of estimation. Treating all steps of the analysis (preprocessing, imputation, feature selection, modeling) as an atom, and cross-validating configurations not just the final modeling step, *avoids overestimation*. A statistical method for removing the performance optimism (bias) due to trying numerous configurations (bootstrapped bias correction cv) is also *necessary to avoid overestimation*. The use of a feature selection algorithm that scales up to hundreds of thousands of biomarkers, suitable for small sample sizes allows the identification of *multiple statistically equivalent biosignatures* (SES). Lastly, notice that the final model suggested for clinical use is constructed on all samples: *no samples are lost to estimation*. On average, the model fit on all data will be the most predictive. All other models are only built to determine the winning configuration and estimate the its expected performance. This signifies a shift in estimation perspective: it the model-producing method that is evaluated, not a specific model instance.

The implications of the above results to a practitioner in translational research is that they can now automatically perform a large percentage of the typical analyses that are relevant to the field. Such analysis may involve multi-omics data complimented with clinical, epidemiological, and lifestyle factors resulting in tens of thousands of measured quantities. This is the case even when sample size is low (< 40); while there is large variance in the guarantees of predictive performance, the user can gauge its clinical utility by the confidence intervals reported. There is no statistical knowledge or expertise required, or computer programming abilities. However, some basic statistical knowledge is still necessary to fully interpret all reported metrics and graphs. Emphasis is placed on facilitating biological interpretation and clinical translation, not only on obtaining a highly predictive model. The results are complemented by (i) confidence intervals of performance to convey clinical utility, (ii) ROC curves that allow one to choose the optimal trade-off between false positive and false negatives for clinical operational use, (iii) visualization of all identified signatures and biomarkers to understand the underlying biology of design diagnostic assays, (iv) metrics of marker impact to the predictive power (importance weighting) to optimize the cost-benefit trade-off of including all or some biomarkers in the model, (iv) a visualization of the role of each marker to the prediction (Individual Conditional Expectation or ICE) plot to facilitate biological interpretation, (v) scatter plots of predictions to identify possibly mislabeled data, (vi) interpretable models that trade-off predictive performance for human interpretability.

The presented use case on Thymic Epithelial Tumors (TET), illustrates the type and breadth of results a practitioner can expect from modern AutoML technologies. TET are relatively rare, still they represent a significant clinical entity with unmet needs in terms of prognosis and therapy. They present a wide phenotypic spectrum broadly divided into two categories, thymomas often associated with autoimmune disorders (in particular thymoma-associated myasthenia gravis) and the less common thymic carcinomas that entail a more aggressive clinical course. **Thus, TET represent a typical biomarker discovery target pathology.** Traditional histological classification on the basis of neoplastic epithelial cell morphology, the degree of the lymphocytic component and the presence or absence of epithelial cell atypia recognizes five subtypes, namely A, AB, B1, B2 and B3, shown to correlate with clinical outcomes^41^. However, histological subtyping has been challenging because of histological complexity, inter-observer inconsistency and lack of prognostic consistency in this morphologically and molecularly heterogeneous group.

In an attempt to identify the genomic underpinnings of TET, Radovich et al. (2018)^27^ used multiplatform omics analyses on 117 TETs. By the combined consideration of thousands of molecular measurements, four distinct molecular subtypes were defined not identical to the classical histological subtypes (A, AB, B, and TC). The centroids of platform-specific cluster assignments from the CNV, mRNA, miRNA, DNA methylation, and reverse phase protein array (RPPA) data were used by a modification of the cluster-of-clusters-assignments approach. This integrated genomic analysis demonstrated that TET are distinct biological entities that do not represent a histological continuum of diseases. We used some of the same datasets and analyzed them through JAD Bio.

After automatically fitting 20890 models, the system found one with almost perfect classification power (AUC = 0.982) to new samples. The fact that just 2 out of the 22796 measured biomarkers carry all information to almost perfectly classify the 4 sub-types of TET **is a surprising yet impressive result**. In addition, there are 45 alternatives of such 2 biomarker signatures. **This multiplicity is particularly favorable to diagnostics, as multiple choices are provided for the future designing a TET subtyping assay.** Furthermore, the set of signatures also conveys a more complete picture of the underlying biology involved. The fact that the second biomarker element (the expression of the CD3E gene) is ranked in the 190^th^ place as ordered by its (unconditional) p-value, demonstrates the utility of modern feature selection algorithms that filter out redundant markers and identify markers that predict *in synergy not in isolation*. This is expected to be the case when markers are involved in independent pathways or pathophysiologically distant molecular events. The example illustrates the difference of signature identification (feature selection) from standard differential expression analysis. The exact molecular pathways that the two molecules underpinned here are involved are subject of further studies, expected to contribute significantly to unfolding underlying biology and inform targeted drug discovery of this complex pathology. **The use of JADBio results to lead data-driven investigations is highlighted.** Interestingly, one of the two identified features, CD3E, is a specific marker of T cell-receptor induced autophagy, considered to be one of the main mechanisms of antigen-presentation in thymus and the molecular basis of autoimmunity^42^. On the other hand, miR-498 has been implicated in many cellular processes, whereas malignant tumors often express low miR-498 levels^43–47^.

There are numerous limitations in the study. The scope of automation and experimental validation concerns low-sample, high-dimensional transcriptomics, methylomics and metabolomics datasets and binary outcomes. Transfer of the conclusions to other types of molecular data (e.g., proteomics, single cell) and outcome types needs further study. The automation architecture presented does not include preprocessing steps of the raw molecular data and signal. It also does not encompass the full range of modern machine learning tasks, such as clustering, representation learning, or causal discovery, to name a few. Predictor types do not include medical images and signals, free medical text, or measurements over time. Prior medical knowledge in the form of biological and medical ontologies or pathways is not considered during the analysis. Datasets are analyzed independently of all other datasets publicly available; recent research directions try to consider datasets in their totality^48^ instead. Biosignatures identification and predictive modeling are but the first steps of a long process in bringing new diagnostic or prognostic tests to the clinic. Follow up steps include, but are not limited to, independent validation studies, the development of tests suitable for the clinical practice, and cost-benefit analyses.

## Methods

### Datasets and Preprocessing

#### The Thymoma dataset

processed data were obtained from the Tumor Molecular Pathology Analysis Working group of the NCI Center for Cancer Genomics Genomic. Molecular features include 2156 binary variables indicating the presence of mutation in their corresponding genes, methylation levels of 3061 CpG sites, 1305 copy number variation measurements, and the expression of 822 miRNAs and 20531 genes (total: 27796 features). Thymoma subclasses were obtained as defined in the original publication^27^. There are 46, 36, 21, and 14 samples to subtypes 1, 2, 3, and 4, respectively.

#### Datasets

we collected data from repositories offering datasets with case-control binary outcomes and include both molecular profiles and curated meta-data (i.e., study design information). BioDataome^32^ is such an online repository with *transcriptome* (both *microarray* and *RNA-seq*) and *epigenetics* (methylation array) datasets. BioDataome uniformly processes and automatically annotates datasets from the Gene Expression Omnibus database (GEO)^1^ and the RECOUNT database^2^. It uses a text-mining pipeline for automatically separating profiles in controls and cases, if applicable, obtaining a dichotomous target for prediction. This automatically assigned status (cases vs. controls) was chosen as the binary outcome of interest.

Metabolomics datasets were obtained from the Metabolomics Workbench^3^, a repository funded by the “NIH Common Fund Metabolomics” initiative. We manually identified a suitable dichotomous prediction target for each metabolomics dataset out of their respective meta-data. As inclusion/exclusion criteria we selected all studies from BioDataome and Metabolomics Workbench with at least 40 samples and for which a binary outcome could be identified. We also reuire at least 10 samples for each class.

For BioDataome we focused on the GPL570 (Affymetrix Human Genome U133 Plus 2.0 Array) and GPL13534 (human methylation 450k beadchip array) and discarded the remaining microarray platforms. These selection criteria lead to the selection of 271 transcriptomics microarray datasets (54675 measurements each, 27510 samples in total), 30 methylation datasets (485512 measurements each, 3546 samples in total), 27 Rna-seq datasets (48674 measurement on average (sd 7585) and 3070 samples in total), and 46 metabolomics datasets (1815 measurement on average (sd 6473) and 4775 samples in total).

Data normalization procedures are detailed in the Supplementary Methods, along with a complete list of the used datasets (Supplementary Table 2).

### The JADBIO architecture

JADBIO has the following functionality and properties: (a) given a 2D matrix of data, it automatically produces predictive models for a discrete (classification), continuous (regression), or time-to-event (survival analysis) outcome. No selection of appropriate algorithms to apply is necessary or tuning of their hyper-parameter values. (b) It identifies multiple equivalent biosignatures, (c) it produces *conservative* predictive performance estimates and corresponding confidence intervals, and (d) it facilitates visualization, explanation, and interpretation of results. In terms of scope and applicability, the current version accepts up to hundreds of thousands of features, and sample sizes as low as a couple of dozen. It accepts numerical measurements (such as omics data), as well as discrete predictors (such as experimental factors, phenotypic and lifestyle predictors), and incomplete datasets with missing values. JADBIO appropriately treats *clustered data* (not to be confused with clustering of data), which are defined as “data that can be classified into a number of distinct groups or ‘clusters’ within a particular study^49^. It is common in this case that samples may still be correlated given the features (or the model). Clustered data violate the most common assumption made by most learning algorithms: that samples are identically and independently distributed (*i.i.d. data*). Examples of clustered data in bioinformatics include matched case-control data, genetic studies including same family members, and repeated measurements on the same subject. JADBIO accepts user-preferences regarding the execution time of the analysis, interpretability of the final model, and enforcing feature selection or not. Regarding efficiency, JADBIO includes numerous optimizations on the algorithms, caching of intermediate results (e.g., in feature selection), and running analysis in parallel, currently on Amazon Web Services (AWS) (https:aws.amazon.com). Finally, JADBIO employs a recently developed^12^ protocol, namely Bootstrap Bias Corrected CV (BBC-CV), for tuning the hyper-parameters of algorithms while estimating performance and adjusting for multiple tries (more details below). The protocol is an order of magnitude faster than the previous state-of-the-art protocol of nested cross validation^11^ and up to two orders of magnitude when faster when heuristics such as the Early Dropping are employed^12^.

JADBIO’s architecture is shown in Figure 6. Two important concepts are the analysis **scenario** and the pipeline **configuration**. A scenario is comprised of all the statistical characteristics of the data and the machine learning task (called meta-level features^50^), as well as the user preferences. These are all aspects that may affect decisions about which algorithms to employ, what are the allowed combinations of algorithms, and what is the space of their hyper-parameter values to explore. An example of a scenario is the analysis of a dataset with ‘tiny’ sample size (<30), ‘huge’ dimensionality (> 100000), binary outcome, no clustered data, for a user that would like to enforce feature selection. A biological such example would be to learn to predict lung cancer from case-control gene expression data.

**Figure 6.**
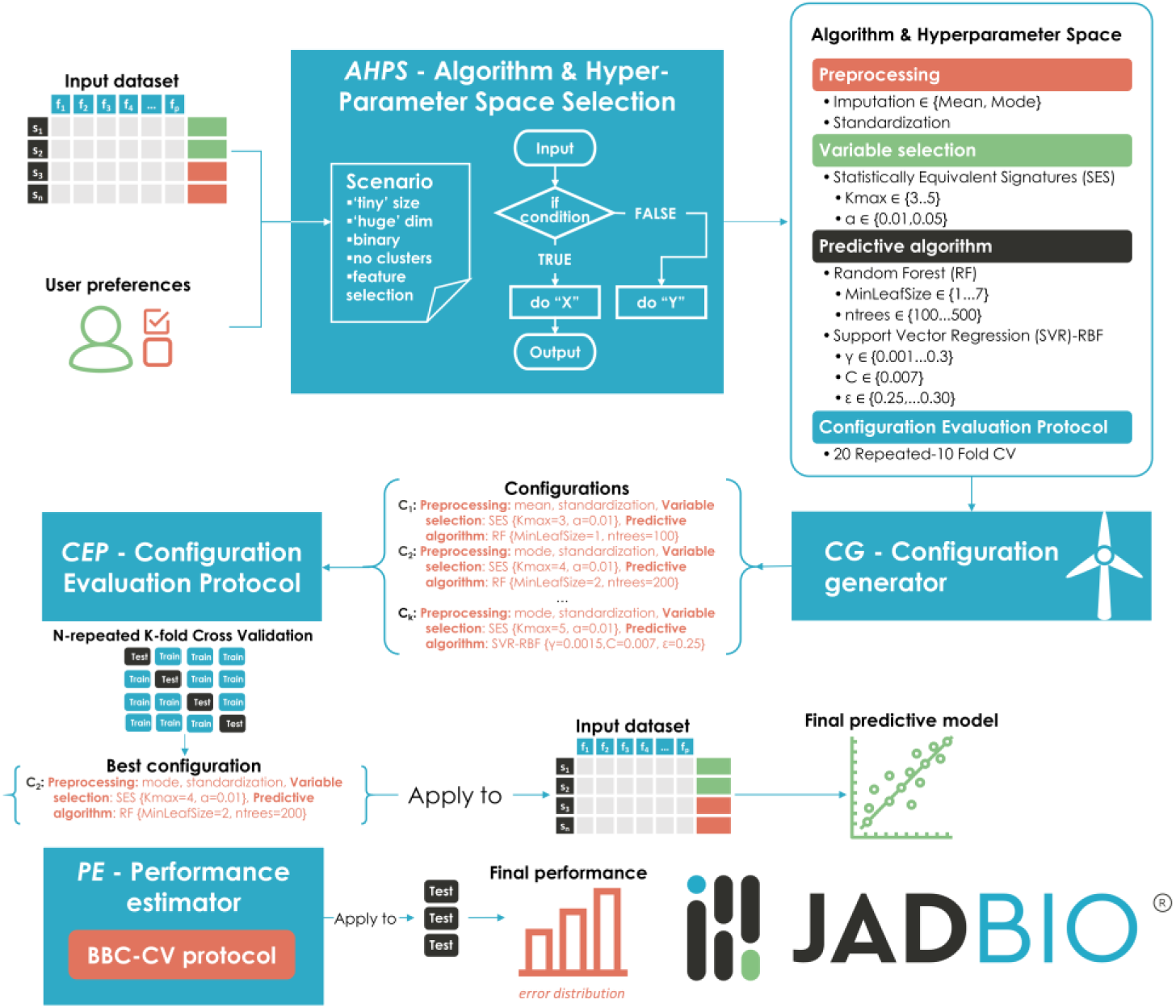
JAD Bio Architecture. The artificial intelligence (AI) engine, namely Algorithm and Hyper-Parameter Space selection *AHPS*, analyzes the current scenario as well as the user’s preferences and selects the most suitable algorithms and hyper-parameter ranges to try, as well as the Configuration Evaluation Protocol (CEP) that estimates performance for each configuration. Using this information, the Configuration Generator (CG) creates the configurations to be evaluated. The winning configuration is applied on all input data (no samples are lost to estimation) to generate the final predictive model. The Performance Estimation protocol is applied on the out-of-sample predictions (indicated by Test boxes) during the CEP to remove the bias due to multiple tries and output the final performance estimate.

A configuration is an instantiation of these choices for each step of the analysis: which combination of algorithms to run and their hyper-parameter values. A configuration results in a pipeline of machine learning steps that accepts the data and outputs a predictive model and estimates of its performance. An example of a configuration is ‘impute missing values with their mean and mode, run the SES algorithm for feature selection with hyper-parameter values *a = 0.05*, and *maxk = 3*, then run a Support Vector Machine with linear kernel and cost hyper-parameter *C = 100”*.

JADBIO contains an Artificial Intelligence (AI) Decision Support System called Algorithm and Hyper-Parameter Space selection (**AHPS**) that, given the scenario, selects the most suitable algorithms and the set of their hyper-parameter values to try for preprocessing, data transformation, imputation of missing values, feature selection, and modeling. It also selects the post-analysis algorithms for visualizing and presenting the results. *AHPS currently differentiates the analysis among about 7000 scenarios.* Quite importantly AHPS selects which Configuration Evaluation Protocol (**CEP**) to be used to estimate predictive performance and identify the optimal configuration. CEPs are explained in detail below. Next, based on these recommendations, the **Configuration Generator** (***CG***) produces a set of configurations to try within the space of choices output by AHPS. Currently, configurations are produced using a static heuristic and grid search^51^ in the space of hyper-parameter values. Depending on the user-preferences for the tuning effort, more or fewer configurations are tried.

Each configuration is fed to the **Configuration Evaluation Protocol**, or **CEP**, that estimates its expected out-of-sample (generalization) predictive performance. JADBIO uses *stratified, N-repeated, K-fold cross-validation* for relatively small sample sizes, and a stratified hold-out or incomplete cross-validation for large sample sizes^12^. We will explain in more detail the former protocol as it is the one for typical bioinformatics scenarios with small sample sizes. Cross-validation is a standard technique where the sample size is partitioned to *K* folds; a configuration produces a model based on all sample folds but one, and a performance estimate is produced on the held-out fold. The final performance of the configuration is the average over all folds. Stratification (in classification problems) implies that each fold in cross-validation follows approximately the same distribution of classes as the un-partitioned dataset; stratification has been shown to provide more robust results for small sample sizes, than without^18^. *N-*repeated cross-validation implies that the cross-validation procedure runs *N* times with different partitions to folds to reduce the variance in the estimation due to the specific partitioning; again, this has been shown to reduce the uncertainty of estimation and to tighten the confidence intervals^12^. The reduced uncertainty in performance estimation also implies that the true best-performing configuration is selected with higher probability, thus repeating the cross-validation also improves predictive performance. The value of *K* depends on the scenario and mostly the total sample size, but also other factors: one needs to also consider in classification problems the samples per class and the imbalance between classes, and the number of censored outcomes in a time-to-event analysis. The number of repetitions is determined by an adaptative procedure, which tracks the confidence intervals’ shrinkage across repeats and stops when there is no progress achieved by the current repetition.

The **CEP** estimates the out-of-sample predictive performance of each configuration and identifies the winner. *The winning configuration is then applied on the whole dataset to produce the final model* (*Final Model Generator*). Thus, *no samples are lost to estimation*. The justification for this is that most learning algorithms produce better predictive models on average with increasing sample size. Thus, the model produced on all samples is better on average than all models produced during cross-validation with the same configuration. These models only serve for performance estimation. The final model output is a new software object that accepts data from the input-data distribution and outputs a prediction. Thus, it is important to notice, that *the model implements all steps learned during the analysis*; a model *is not* just a Decision Tree or a Support Vector Machine. A model includes the code to normalize features, to impute missing values, and to project on the selected features.

Next, JADBIO estimates the generalization predictive performance of the average model produced by the winning configuration. Unfortunately, **the cross-validated performance of the winning configuration is overestimated and should not be returned**. For example, if the cross-validated performance of the winning configuration is 90%, the true expected performance of this configuration is less than 90% (or equal in degenerate cases). To be more precise, the expected performance of the models produced by the winning configuration when trained on data from the given data distribution and size equal to the training sets during cross-validation, is less than 90%. This is because multiple configurations are tried and thus, some adjustment is necessary; the phenomenon is conceptually equivalent to the adjustment of *p*-values in multiple hypothesis testing. In the context of multiple machine learning models, it is called the “multiple comparison in induction algorithms problem”^52^, and it is related to the “*winner’s curse”* in statistics^53^. Intuitively, when selecting the best performing algorithm among many, it is likely that it is because it achieved a performance higher than its average. The overestimation has been proved theoretically, but also shown empirically in Tsamardinos et al. (2018)^12^ as well as in Figure 4, and is particularly problematic in small-sample analyses like the ones encountered in bioinformatics. JADBIO’s **Performance Estimator** (**PE**) employs the BBC-CV protocol to return slightly conservative performance estimates^12^. The main idea of these protocols is to bootstrap the configuration selection strategy on the out-of-sample prediction matrix produced during the cross-validation. BBC-CV removes the bias due to the multiple tries. It is one order-of-magnitude more efficient than the previous standard protocol, namely the *nested cross-validation*^11^. The BBC-CV protocol can output with no additional computational overhead the probability distribution of the expected performances and their 95% confidence intervals. For classification problems, an adaptation of the BBC-CV idea can also produce *adjusted-for-multiple-tries* ROC curves along with their confidence intervals.

There are two major points to notice, which guarantee the correctness of performance estimation even for tiny sample sizes. The first one is the application of a protocol that adjusts performance for multiple tries, as discussed above. The second regards the fact that *a configuration is crossvalidated as an atom*: the imputation, normalization, standardization, feature selection, and modelling algorithms are jointly cross-validated as one procedure. This avoids a common methodological mistake in omics analyses where preprocessing, normalization, imputation, and feature selection are first applied on *the complete dataset* and only the last step of modelling (e.g., the Decision Tree algorithm) is cross-validated^11^. The error could lead to significant overestimation of performance, overfitting, and invalidation of results: an eye-opening pedagogical experiment is presented at page 245 of Hastie, Tibshirani, and Friedman’s book^54^, where the true error rate is 50% but the estimate is only 3% error if one applies first the feature selection step on all data and only then cross-validates the learning algorithm.

During the analysis JADBIO reports the current best performance over all tried configurations and its confidence interval. This allows the user to stop the analysis early if a desired predictive performance has been reached. Once the analysis is over, several pieces of information are presented, as showcased in the results section on the Thymoma data.

### JADBIO Evaluation Experimentation Protocol

To produce the results in Figures 4-5, JADBIO was applied on a wide palette of datasets (Supplementary Table 2) and its performance was evaluated on independent hold out sets. First, each original dataset was randomly split in half in a stratified way (i.e., maintaining the binary outcome distribution); the JADBIO pipeline was then applied in turn on each split, used as the training set, while the other split is held out for testing purposes (holdout set). In each run JADBIO returns the best model over all possible configurations, an estimation of out-of-sample performance along with its bootstrap-based performance estimates. The JADBIO estimated performance on the training set is then compared against the achieved performance of the returned model on the hold out set. The process is repeated when we enforce feature selection and when not, and when restricting the tried configurations to only the ones that produce humanly interpretable models. To produce the confidence intervals on the hold out test sets, we use 1000 bootstraps.

### Comparative Evaluation Protocol Against auto-sklearn

We present a comparative evaluation of auto-sklearn against JADBIO on the classification analyses. Specifically, for each run, the auto-sklearn library was applied on the same training and holdout sets used by JADBIO, using the same computational architecture and number of CPUs. The maximum time limit for auto-sklearn was set to the maximum between one hour and the termination time of JADBIO; runs that exceeded twice this limit were forcefully terminated. The same performance metric, namely AUC, was employed by both auto-sklearn and JADBIO for identifying the best performing model. All other auto-sklearn hyper-parameters were set to their default values. To note, the library does not allow to identify which variables are included in the final model. Additional information on the inner operation of auto-sklearn are available in the Supplementary Methods.

### Data availability and software notes

JADBIO is available at http://jadbio.com, and can be used for reproducing all results. A free trial version is offered. All the data used in the analysis are freely available at BioDataome (http://dataome.mensxmachina.org/) and Metabolomics Workbench (https://www.metabolomicsworkbench.org/).

## Acknowledgments

We would like to thank the TCGA consortium for providing us with the thymoma data.

## Author contributions

IT conceived the study and designed the algorithms. VL performed and supervised the study. IT and VL wrote the main part of the manuscript. PC and GB significantly contributed to the implementation of JADBIO. JCZ provided the thymoma case and interpretations. KL helped with the figures. All co-authors contributed to the writing of the manuscript.

## References

1. Barrett, T. et al. NCBI GEO: Archive for functional genomics data sets - Update. Nucleic Acids Res. 41, (2013).

2. Collado-Torres, L. et al. Reproducible RNA-seq analysis using recount2. Nat. Biotechnol. 35, 319–321 (2017).

3. Sud, M. et al. Metabolomics Workbench: An international repository for metabolomics data and metadata, metabolite standards, protocols, tutorials and training, and analysis tools. Nucleic Acids Res. 44, D463–70 (2016).

4. Grossman, R. L. et al. Toward a shared vision for cancer genomic data. New England Journal of Medicine (2016). doi:10.1056/NEJMp1607591

5. Teschendorff, A. E. Avoiding common pitfalls in machine learning omic data science. Nat. Mater. 18, 422–427 (2019).

6. Feurer, M., Eggensperger, K., Falkner, S., Lindauer, M. & Hutter, F. Practical Automated Machine Learning for the AutoML Challenge 2018. in International Workshop on Automatic Machine Learning at ICML (2018).

7. Johnstone, I. M. & Titterington, D. M. Statistical challenges of high-dimensional data. Philos. Trans. R. Soc. A Math. Phys. Eng. Sci. (2009). doi:10.1098/rsta.2009.0159

8. Ioannidis, J. P. Microarrays and molecular research: noise discovery? Lancet 365, 454–455 (2005).

9. Michiels, S., Koscielny, S. & Hill, C. Prediction of cancer outcome with microarrays: A multiple random validation strategy. Lancet (2005). doi:10.1016/S0140-6736(05)17866-0

10. Tsamardinos, I., Aliferis, C. F. & Statnikov, A. Time and sample efficient discovery of Markov blankets and direct causal relations. in Proceedings of the ACM SIGKDD International Conference on Knowledge Discovery and Data Mining (2003). doi:10.1145/956750.956838

11. Statnikov, A., Aliferis, C. F., Tsamardinos, I., Hardin, D. & Levy, S. A comprehensive evaluation of multicategory classification methods for microarray gene expression cancer diagnosis. Bioinformatics 21, 631–643 (2005).

12. Tsamardinos, I., Greasidou, E. & Borboudakis, G. Bootstrapping the out-of-sample predictions for efficient and accurate cross-validation. Mach. Learn. 107, 1895–1922 (2018).

13. Statnikov, A., Tsamardinos, I., Dosbayev, Y. & Aliferis, C. F. GEMS: a system for automated cancer diagnosis and biomarker discovery from microarray gene expression data. Int. J. Med. Inform. 74, 491–503 (2005).

14. Tsamardinos, I., Brown, L. E. & Aliferis, C. F. The max-min hill-climbing Bayesian network structure learning algorithm. Mach. Learn. 65, 99–111 (2006).

15. Aliferis, C. F., Statnikov, A. R., Tsamardinos, I., Mani, S. & Koutsoukos, X. D. Local Causal and Markov Blanket Induction for Causal Discovery and Feature Selection for Classification Part I : Algorithms and Empirical Evaluation. J. Mach. Learn. Res. 11, 171–234 (2010).

16. Lagani, V. & Tsamardinos, I. Structure-based variable selection for survival data. Bioinformatics 26, 1887–1894 (2010).

17. Lagani, V., Kortas, G. & Tsamardinos, I. Biomarker signature identification in ‘omics’ data with multi-class outcome. Comput. Struct. Biotechnol. J. 6, (2013).

18. Tsamardinos, I., Rakhshani, A. & Lagani, V. Performance-estimation properties of cross-validation-based protocols with simultaneous hyper-parameter optimization. in Lecture Notes in Computer Science 8445 LNCS, 1–14 (2014).

19. Tsagris, M., Lagani, V. & Tsamardinos, I. Feature selection for high-dimensional temporal data. BMC Bioinformatics 19, (2018).

20. Lagani, V., Athineou, G., Farcomeni, A., Tsagris, M. & Tsamardinos, I. Feature Selection with the R Package MXM: Discovering Statistically-Equivalent Feature Subsets. J. Stat. Softw. 80, (2017).

21. Borboudakis, G. et al. Chemically intuited, large-scale screening of MOFs by machine learning techniques. npj Comput. Mater. 3, 40 (2017).

22. 22. Adamou, M. et al. Toward Automatic Risk Assessment to Support Suicide Prevention. Crisis 40, 249–256 (2019).

23. Simantiraki, O., Charonyktakis, P., Pampouchidou, A., Tsiknakis, M. & Cooke, M. Glottal source features for automatic speech-based depression assessment. in Proceedings of the 18th Conference of the International Speech Communication Association INTERSPEECH 2700–2704 (2017). doi:10.21437/Interspeech.2017-1251

24. Agrapetidou, A., Charonyktakis, P., Gogas, P., Papadimitriou, T. & Tsamardinos, I. An AutoML application to forecasting bank failures. Appl. Econ. Lett. 1–5 (2020). doi:10.1080/13504851.2020.1725230

25. Orfanoudaki, G., Markaki, M., Chatzi, K., Tsamardinos, I. & Economou, A. MatureP: prediction of secreted proteins with exclusive information from their mature regions. Sci. Rep. 7, 3263 (2017).

26. Panagopoulou, M. et al. Circulating cell-free DNA in breast cancer: size profiling, levels, and methylation patterns lead to prognostic and predictive classifiers. Oncogene 38, 3387–3401 (2019).

27. Radovich, M. et al. The Integrated Genomic Landscape of Thymic Epithelial Tumors. Cancer Cell (2018). doi:10.1016/j.ccell.2018.01.003

28. Breiman, L. Random Forests. Mach. Learn. 45, 5–32 (2001).

29. Tsamardinos, I. & Aliferis, C. F. Towards principled feature selection: relevancy, filters, and wrappers. in Proceedings of the Ninth International Workshop on Artificial Intelligence and Statistics (2003).

30. Statnikov, A. & Aliferis, C. F. Analysis and computational dissection of molecular signature multiplicity. PLoS Comput. Biol. (2010). doi:10.1371/journal.pcbi.1000790

31. Goldstein, A., Kapelner, A., Bleich, J. & Pitkin, E. Peeking Inside the Black Box: Visualizing Statistical Learning With Plots of Individual Conditional Expectation. J. Comput. Graph. Stat. 24, 44–65 (2015).

32. Lakiotaki, K., Vorniotakis, N., Tsagris, M., Georgakopoulos, G. & Tsamardinos, I. BioDataome: a collection of uniformly preprocessed and automatically annotated datasets for data-driven biology. Database 2018, (2018).

33. Cleveland, W. S., Grosse, E. & Shyu, W. M. Local Regression Models. in Statistical Models in S (eds. Chambers, J. M. & Hastie, T. J.) (Chapman and Hall, 2017). doi:10.1201/9780203738535

34. Domingos, P. & Domingos, P.A Unified Bias-Variance Decomposition and its Applications. PROC. 17TH Int. CONF. Mach. Learn. 231–238 (2000).

35. Do we Need Hundreds of Classifiers to Solve Real World Classification Problems? J. Mach. Learn. Res. 15, 3133–3181 (2014).

36. Feurer, M. et al. Efficient and Robust Automated Machine Learning. in Advances in Neural Information Processing Systems 28 (eds. Cortes, C., Lawrence, N. D., Lee, D. D., Sugiyama, M. & Garnett, R.) 2962–2970 (Curran Associates, Inc., 2015).

37. ChaLearn. AutoML.

38. Xanthopoulos, I., Tsamardinos, I., Christophides, V., Simon, E. & Salinger, A. Putting the human back in the AutoML loop. in CEUR Workshop Proceedings (2020).

39. Thornton, C., Hutter, F., Hoos, H. H. & Leyton-Brown, K. Auto-WEKA: Combined selection and hyperparameter optimization of classification algorithms. in Proceedings of the ACM SIGKDD International Conference on Knowledge Discovery and Data Mining (2013). doi:10.1145/2487575.2487629

40. Aliferis, C. F. et al. Factors influencing the statistical power of complex data analysis protocols for molecular signature development from microarray data. PLoS One (2009). doi:10.1371/journal.pone.0004922

41. Marx, A. et al. ITMIG consensus statement on the use of the WHO Histological classification of thymoma and thymic carcinoma: Refined definitions, histological criteria, and Reporting. J. Thorac. Oncol. (2014). doi:10.1097/JTO.0000000000000154

42. Barbouti, A. et al. Implications of Oxidative Stress and Cellular Senescence in Age-Related Thymus Involution. Oxid. Med. Cell. Longev. (2020). doi:10.1155/2020/7986071

43. Liu, R., Liu, F., Li, L., Sun, M. & Chen, K. MiR-498 regulated FOXO3 expression and inhibited the proliferation of human ovarian cancer cells. Biomed. Pharmacother. (2015). doi:10.1016/j.biopha.2015.04.005

44. Gopalan, V., Smith, R. A. & Lam, A. K. Y. Downregulation of microRNA-498 in colorectal cancers and its cellular effects. Exp. Cell Res. (2015). doi:10.1016/j.yexcr.2014.08.006

45. Islam, F. et al. MiR-498 in esophageal squamous cell carcinoma: clinicopathological impacts and functional interactions. Hum. Pathol. (2017). doi:10.1016/j.humpath.2017.01.014

46. Xiang, H. Y. et al. Upregulation of miR-498 suppresses Th17 cell differentiation by targeting STAT3 in rheumatoid arthritis patients. Sheng Li Xue Bao (2018).

47. Wang, T., Ma, L., Li, W., Ding, L. & Gao, H. MicroRNA-498 reduces the proliferation and invasion of colorectal cancer cells via targeting Bcl-2. FEBS Open Bio (2020). doi:10.1002/2211-5463.12767

48. Lakiotaki, K. et al. A data driven approach reveals disease similarity on a molecular level. npj Syst. Biol. Appl. 5, 1–10 (2019).

49. Galbraith, S., Daniel, J. A. & Vissel, B. A Study of Clustered Data and Approaches to Its Analysis. J. Neurosci. 30, 10601–10608 (2010).

50. Vilalta, R., Giraud-Carrier, C. G., Brazdil, P. & Soares, C. Using Meta-Learning to Support Data Mining. IJCSA (2004).

51. Hsu, C.-W., Chang, C.-C. & Lin, C.-J. A Practical Guide to Support Vector Classification. BJU international (2008).

52. Jensen, D. D. & Cohen, P. R. Multiple comparisons in induction algorithms. Mach. Learn. (2000). doi:10.1023/A:1007631014630

53. Ioannidis, J. P. A. Why most discovered true associations are inflated. Epidemiology (2008). doi:10.1097/EDE.0b013e31818131e7

54. Hastie, T., Tibshirani, R. & Friedman, J. H. The elements of statistical learning : data mining, inference, and prediction. (Springer, 2016).

55. Piccolo, S. R. et al. A single-sample microarray normalization method to facilitate personalized-medicine workflows. Genomics 100, 337–344 (2012).

56. Anders, S. & Huber, W. Differential expression analysis for sequence count data. Genome Biol. 11, R106 (2010).

57. Aryee, M. J. et al. Minfi: a flexible and comprehensive Bioconductor package for the analysis of Infinium DNA methylation microarrays. Bioinformatics 30, 1363–1369 (2014).

58. Du, P. et al. Comparison of Beta-value and M-value methods for quantifying methylation levels by microarray analysis. BMC Bioinformatics 11, 587 (2010).

59. Hutter, F., Hoos, H. H. & Leyton-Brown, K. Sequential model-based optimization for general algorithm configuration. in International Conference on Learning and Intelligent Optimization 507–523 (2011).

60. Pedregosa, F. et al. Scikit-learn: Machine Learning in {P}ython. J. Mach. Learn. Res. 12, 2825–2830 (2011).

