## Supplementary material for "Just Add Data: Automated Predictive Modeling and BioSignature Discovery"

**Abstract**:

Fully automated machine learning, statistical modelling, and artificial intelligence for predictive modeling is becoming a reality, giving rise to the field of Automated Machine Learning (AutoML). AutoML systems promise to democratize data analysis to non-experts, drastically increase productivity, improve replicability of the statistical analysis, facilitate the interpretation of results, and shield against common methodological analysis pitfalls. We present the basic ideas and principles of Just Add Data Bio (JADBIO), an AutoML technology applicable to the low-sample, high-dimensional omics data that arise in translational medicine and bioinformatics applications. In addition to predictive and diagnostic models ready for clinical use, JADBIO also returns the corresponding biosignatures, i.e., minimal-size subsets of biomarkers that are jointly predictive of the outcome of interest. A use-case on thymic epithelial tumors is presented, along with an extensive evaluation on 374 public biological datasets. Results show that long-standing challenges with overfitting and overestimation of complex non-linear machine learning pipelines on high-dimensional, low small sample data can be overcome.

**Supplementary Materials**

### Supplementary Methods

#### *Data normalization procedures*

Normalization of microarray gene expression data was performed in BioDataome using the single-channel array normalization (SCAN) algorithm[55]. SCAN normalizes each array *independently*, ensuring that measurements from profiles in the test sets do not affect the preprocessing of profiles in the training sets; thus, *there is no information leakage from the test sets during cross-validation to the estimation of performance from the training set*. Count values for RNA-seq data were downloaded as prepared by the RECOUNT repository. Each sample was then *independently* normalized for library size (estimated as the sum of all its reads) and log2-transformed. More sophisticated methods do exist both for library size normalization and variance-stabilizing transformation (see for example the approaches provided in the DESeq2 R package[56]), but they do not preprocess samples independently and would require special treatment, i.e., to be incorporated within the cross-validation procedure. Background correction and normalization of methylation data was carried out with the minfi R package[57], with beta values used for all subsequent analyses[58].

#### *Details on auto-sklearn internal operation*

Auto-sklearn is a ML library that automates some ML functionalities. The core of auto-sklearn is the Sequential Model-Based Optimization (SMBO) algorithm[59] for searching the space of configurations in an intelligent way. In brief, SMBO uses Gaussian Processes (GP) for fitting the function F that links configurations to predictive performances. Function F is approximated based on the currently examined configurations; the GP also estimates the uncertainty at each point of the configuration space. The configuration to be tested at each step is chosen by taking into account both the region of maximal performance predicted according to F, and the uncertainty of the GP fitting, thus balancing exploitation and exploration in the search of the configurations space. Configurations are built on top of algorithms in the scikit-learn library[60]. Auto-sklearn pipelines require the user to provide a training dataset along with the maximum time limit for the SMBO operation. Time limits for the SMBO algorithm are not strongly enforced, with the algorithm typically terminating after the deadline. By default, configurations are tested using a hold-out protocol, where 70% of the samples are used for training and 30% held out for assessing performances. The auto-sklearn output is an ensemble model comprising of a subset (by default 50) of the best performing models found by the SMBO algorithm. The library offers limited functionalities for investigating the inner operation of the resulting model; particularly, it is not possible to identify which variables out of the initial ones are employed in the models included in the final ensemble. Consequently, auto-sklearn models always require a dataset including all original predictors in order to provide new predictions.

#### *Thymoma measurements codebook*

All features follow the structure:

data type : platform ID 1 : platform ID 2 : feature ID 1 : feature ID 2 :

- Data type can be binary (B), integer (I), categorical (C), numerical (N).
- The platform ID 1 indicates the type of omics data. platform used. DNA mutation calls (MUTA), DNA methylation beta values (METH), mRNA gene expression levels (MRNA), miRNA expression levels (MIR), copy number variation scores (CNVR).
- The platform ID 2 indicate specific type of data within an omics platform, and can be empty. For example, HOTS is used to denote hotspot mutations.
- The feature ID 1 is the name of the feature, typically the symbol associated to the gene.
- The feature ID 2 helps in better identifying the feature if necessary, and can be empty.

*Methods for post-analysis explanation, visualization, and model interpretation*

The results produced by JADBIO platform are presented to the user using different graphical representations.

Individual Variable Importance Plots: The purpose of this plot is to assess the added value of each selected feature. Individual variable importance measures the effect in predictive information when a single feature (variable) is removed. For each variable in turn, individual variable importance is computed as the ratio between the resulting cross-validated performance when the variable is removed and the cross-validated performance obtained on the original dataset; in both cases, the winning configuration is employed to build the models. To be more precise, a variable is not removed from the data, but its values are permuted instead, thus ensuring that it carries no predictive information. Virtually removing the variable through permutation, instead of removing it completely during cross-validation ensures that the dimensionality of the problem remains the same in both cases and that the best identified hyper-parameter values for all algorithms do not need to be adjusted for differences in the dimensionality of the learning task. The permutation technique is conceptually similar to the calculation of importance weighting in Random Forests [28]. An example on the Thymoma use-case is in **Σφάλμα! Το αρχείο προέλευσης της αναφοράς δεν βρέθηκε.**d (left-bottom).

Individual Conditional Expectation (ICE) plots. ICE plots visualize the effect of the value of a given variable to the prediction of the model. Specifically, for each variable $X$ (e.g., a specific gene expression), the value $f(X = x)$ is plotted, where $f$ is the prediction of the model, e.g,. the probability of being a disease sample. Since the actual prediction depends not only on a single variable, but all the variables selected in the model, the ICE plots the average $f(X = x)$ over the empirical distribution of the value combinations of the variables, along with their confidence intervals. The method is described in detail by Goldstein and co-authors[31]. The ICE plot conveys useful information. The user can gauge whether observing an increased value of a variable *X* results in higher or lower probability of disease (for disease outcomes, of course). It is possible that the ICE curves are not monotonic, e.g., as drug dosage increases, the probability of successful treatment increases and the decreases again, if the drug becomes toxic. Also, the variance (confidence intervals) of the plot are quite informative: values of *X* with low variance in the prediction imply that the model does not consider much the values of the other variables. For values of *X* with high variance, mean that the model relies more on the other variables to make predictions.

### Supplementary Figures

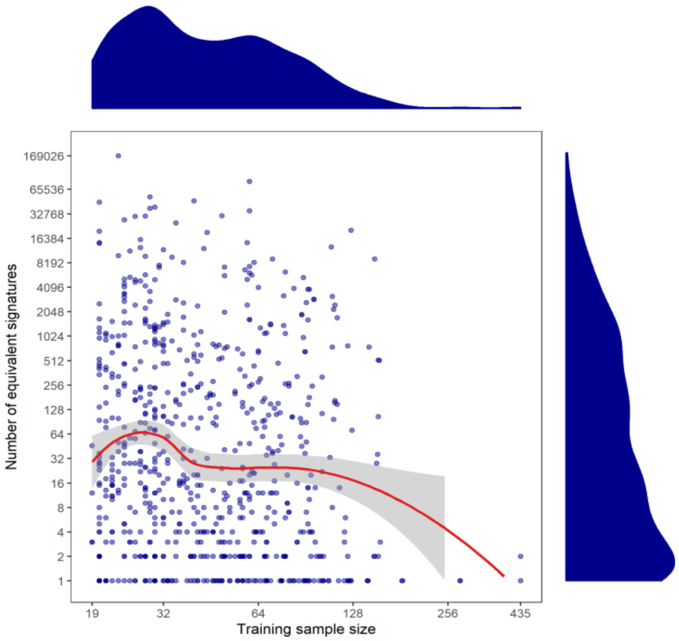

**Supplementary Figure 1: Relationship between signature multiplicity and sample size.** Sample size on the x-axis, number of equivalent signatures on the y-axis. Loess regression depicted as red line, with 95% confidence interval denoted by grey area. There is no apparent trend in the interval 20 to 128 samples, with a sharp decline in number of equivalent signatures only for the runs with more than 200 samples.

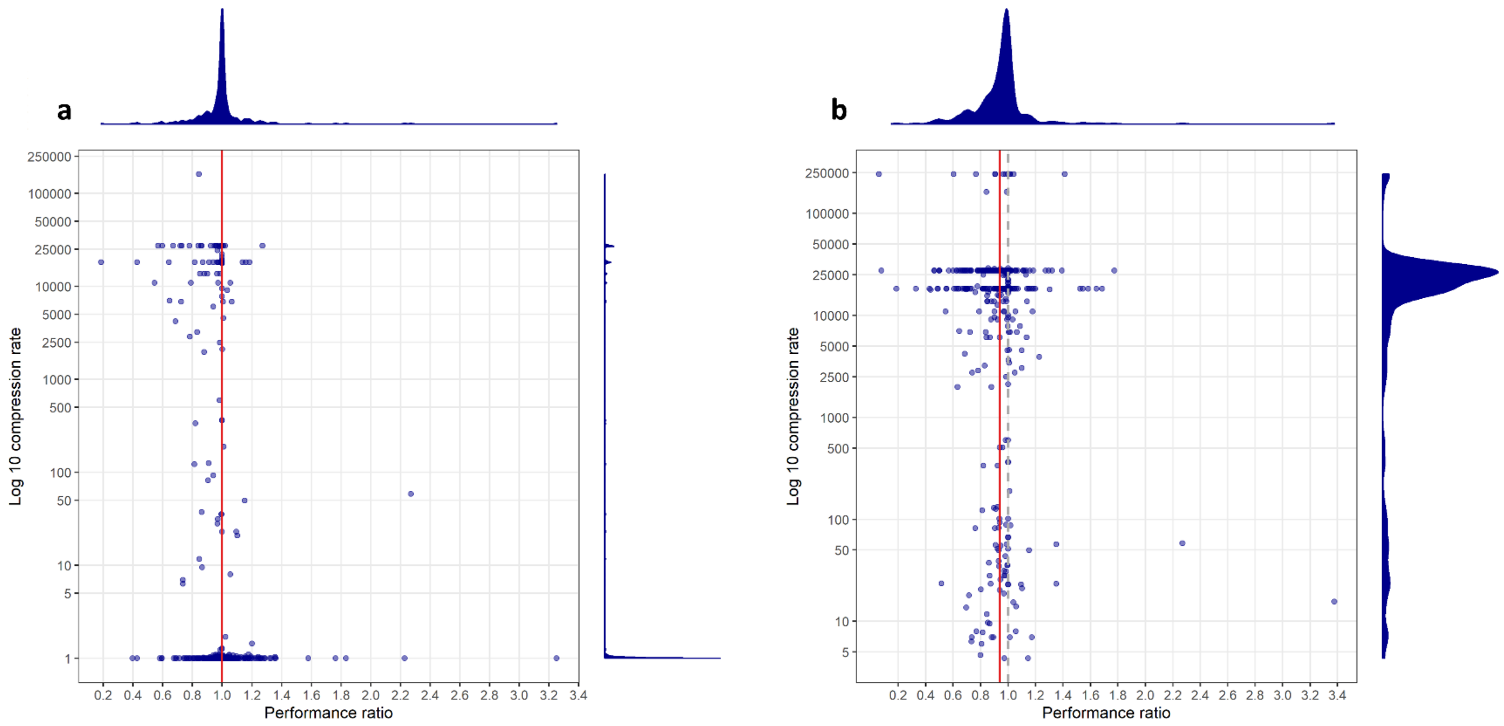

**Supplementary Figure 2: Contrasting JADBIO and auto-sklearn**. Results on the 453 runs successfully completed by auto-sklearn. In both panels each dot represents one run. The x-axis reports the ratio between the holdout AUC achieved by JADBIO over the one achieved by auto-sklearn. The more a point is shifted on the right, the better the performance of JADBIO versus auto-sklearn in the corresponding run. The y-axis reports the ratio between the number of predictors used by auto-sklearn (always corresponding to all measurements) over the number of variables selected by JADBIO. The higher the point, the more parsimonious the model produced by JADBIO. The red vertical line corresponds to the mean value in terms of performance ratio, while the dotted, grey vertical line marks the case of equal performances, i.e., performance ratio equal to 1. A) JADBIO model search is not restricted over the configurations to employ. Notably, the two systems have on average the same performances, however in 23.52% of the runs JADBIO uses more parsimonious (i.e., employing fewer variables) models. B) JADBIO model search is restricted to configurations with feature selection. In this case JADBIO achieves on average 94% of the performances of auto-sklearn, however by constantly using fewer than 25 predictors.

### Supplementary Tables

**Supplementary Table 1: Equivalent signatures identified by JADBIO on the Thymoma dataset.**

| Predictor 1 (miRNA expression) | Predictor 2 (gene expression) |
|---|---|
| N:MIR::hsa-miR-498:Uncorrected: | N:GEXP::CD3E:916: |
| N:MIR::hsa-miR-498:Uncorrected: | N:GEXP::TCF7:6932: |
| N:MIR::hsa-miR-498:Uncorrected: | N:GEXP::GRAP2:9402: |
| N:MIR::hsa-miR-498:Uncorrected: | N:GEXP::TRAF3IP3:80342: |
| N:MIR::hsa-miR-498:Uncorrected: | N:GEXP::ZAP70:7535: |
| N:MIR::hsa-miR-498:Uncorrected: | N:GEXP::CD247:919: |
| N:MIR::hsa-miR-498:Uncorrected: | N:GEXP::LEF1:51176: |
| N:MIR::hsa-miR-498:Uncorrected: | N:GEXP::ARHGDIB:397: |
| N:MIR::hsa-miR-498:Uncorrected: | N:GEXP::SASH3:54440: |
| N:MIR::hsa-miR-498:Uncorrected: | N:GEXP::ARHGAP15:55843: |
| N:MIR::hsa-miR-498:Uncorrected: | N:GEXP::SPN:6693: |
| N:MIR::hsa-miR-498:Uncorrected: | N:GEXP::SH2D1A:4068: |
| N:MIR::hsa-miR-498:Uncorrected: | N:GEXP::CD3G:917: |
| N:MIR::hsa-miR-498:Uncorrected: | N:GEXP::PAFAH2:5051: |
| N:MIR::hsa-miR-498:Uncorrected: | N:GEXP::TMEM71:137835: |
| N:MIR::hsa-miR-517-5p:Uncorrected: | N:GEXP::CD3E:916: |
| N:MIR::hsa-miR-517-5p:Uncorrected: | N:GEXP::TCF7:6932: |
| N:MIR::hsa-miR-517-5p:Uncorrected: | N:GEXP::GRAP2:9402: |
| N:MIR::hsa-miR-517-5p:Uncorrected: | N:GEXP::TRAF3IP3:80342: |
| N:MIR::hsa-miR-517-5p:Uncorrected: | N:GEXP::ZAP70:7535: |
| N:MIR::hsa-miR-517-5p:Uncorrected: | N:GEXP::CD247:919: |
| N:MIR::hsa-miR-517-5p:Uncorrected: | N:GEXP::LEF1:51176: |
| N:MIR::hsa-miR-517-5p:Uncorrected: | N:GEXP::ARHGDIB:397: |
| N:MIR::hsa-miR-517-5p:Uncorrected: | N:GEXP::SASH3:54440: |
| N:MIR::hsa-miR-517-5p:Uncorrected: | N:GEXP::ARHGAP15:55843: |
| N:MIR::hsa-miR-517-5p:Uncorrected: | N:GEXP::SPN:6693: |
| N:MIR::hsa-miR-517-5p:Uncorrected: | N:GEXP::SH2D1A:4068: |
| N:MIR::hsa-miR-517-5p:Uncorrected: | N:GEXP::CD3G:917: |
| N:MIR::hsa-miR-517-5p:Uncorrected: | N:GEXP::PAFAH2:5051: |
| N:MIR::hsa-miR-517-5p:Uncorrected: | N:GEXP::TMEM71:137835: |
| N:MIR::hsa-miR-517c-3p:Uncorrected: | N:GEXP::CD3E:916: |
| N:MIR::hsa-miR-517c-3p:Uncorrected: | N:GEXP::TCF7:6932: |
| N:MIR::hsa-miR-517c-3p:Uncorrected: | N:GEXP::GRAP2:9402: |
| N:MIR::hsa-miR-517c-3p:Uncorrected: | N:GEXP::TRAF3IP3:80342: |
| N:MIR::hsa-miR-517c-3p:Uncorrected: | N:GEXP::ZAP70:7535: |
| N:MIR::hsa-miR-517c-3p:Uncorrected: | N:GEXP::CD247:919: |
| N:MIR::hsa-miR-517c-3p:Uncorrected: | N:GEXP::LEF1:51176: |
| N:MIR::hsa-miR-517c-3p:Uncorrected: | N:GEXP::ARHGDIB:397: |

| | |
|---|---|
| N:MIR::hsa-miR-517c-3p:Uncorrected: | N:GEXP::SASH3:54440: |
| N:MIR::hsa-miR-517c-3p:Uncorrected: | N:GEXP::ARHGAP15:55843: |
| N:MIR::hsa-miR-517c-3p:Uncorrected: | N:GEXP::SPN:6693: |
| N:MIR::hsa-miR-517c-3p:Uncorrected: | N:GEXP::SH2D1A:4068: |
| N:MIR::hsa-miR-517c-3p:Uncorrected: | N:GEXP::CD3G:917: |
| N:MIR::hsa-miR-517c-3p:Uncorrected: | N:GEXP::PAFAH2:5051: |
| N:MIR::hsa-miR-517c-3p:Uncorrected: | N:GEXP::TMEM71:137835: |

**Supplementary Table 2: Number and modality type of datasets employed in the evaluation.**

| | |
|---|---|
| Metabolomics | 46 |
| Transcriptomics (RNA-seq) | 27 |
| Transcriptomics (microarray) | 271 |
| Methylation (microarray) | 30 |
| Total | 374 |

**Supplementary Table 3: Complete list of datasets**. These are the datasets involved in JADBIO large scale evaluation and comparison against auto-sklearn. For each dataset we report from left to right: its identifier, omics data type, number of variables, number of samples, the repository from which it originates, and for annotated datasets also the disease and the generic disease type.

| Name | Type | # variables | # samples | Origin | Disease | Disease type |
|---|---|---|---|---|---|---|
| ST000005 | metabolomics | 56 | 87 | Metabolomics Workbench | | |
| ST000009 | metabolomics | 647 | 114 | Metabolomics Workbench | | |
| ST000011 | metabolomics | 303 | 40 | Metabolomics Workbench | | |
| ST000014 | metabolomics | 132 | 72 | Metabolomics Workbench | | |
| ST000015 | metabolomics | 55 | 125 | Metabolomics Workbench | | |
| ST000016 | metabolomics | 1194 | 110 | Metabolomics Workbench | | |
| ST000017 | metabolomics | 1009 | 42 | Metabolomics Workbench | | |
| ST000062 | metabolomics | 153 | 97 | Metabolomics Workbench | | |
| ST000063 | metabolomics | 167 | 94 | Metabolomics Workbench | | |
| ST000105 | metabolomics | 755 | 111 | Metabolomics Workbench | | |
| ST000106 | metabolomics | 1010 | 64 | Metabolomics Workbench | | |
| ST000135 | metabolomics | 17 | 58 | Metabolomics Workbench | | |
| ST000138 | metabolomics | 12 | 367 | Metabolomics Workbench | | |
| ST000163 | metabolomics | 35524 | 100 | Metabolomics Workbench | | |

| | | | | |
|---|---|---|---|---|
| ST000168 | metabo-lomics | 38 | 200 | Metabolomics Work-bench |
| ST000191 | metabo-lomics | 13 | 56 | Metabolomics Work-bench |
| ST000195 | metabo-lomics | 26 | 51 | Metabolomics Work-bench |
| ST000196 | metabo-lomics | 28 | 135 | Metabolomics Work-bench |
| ST000209 | metabo-lomics | 20 | 48 | Metabolomics Work-bench |
| ST000212 | metabo-lomics | 20 | 60 | Metabolomics Work-bench |
| ST000244 | metabo-lomics | 46 | 51 | Metabolomics Work-bench |
| ST000252 | metabo-lomics | 80 | 40 | Metabolomics Work-bench |
| ST000286 | metabo-lomics | 366 | 40 | Metabolomics Work-bench |
| ST000336 | metabo-lomics | 45 | 58 | Metabolomics Work-bench |
| ST000353 | metabo-lomics | 93 | 42 | Metabolomics Work-bench |
| ST000355 | metabo-lomics | 129 | 214 | Metabolomics Work-bench |
| ST000356 | metabo-lomics | 102 | 134 | Metabolomics Work-bench |
| ST000393 | metabo-lomics | 70 | 68 | Metabolomics Work-bench |
| ST000396 | metabo-lomics | 137 | 299 | Metabolomics Work-bench |
| ST000401 | metabo-lomics | 164 | 40 | Metabolomics Work-bench |
| ST000402 | metabo-lomics | 116 | 71 | Metabolomics Work-bench |

| | | | | | | |
|---|---|---|---|---|---|---|
| ST000421 | metabo-lomics | 20126 | 56 | Metabolomics Work-bench | | |
| ST000422 | metabo-lomics | 19088 | 60 | Metabolomics Work-bench | | |
| ST000450 | metabo-lomics | 245 | 84 | Metabolomics Work-bench | | |
| ST000477 | metabo-lomics | 148 | 113 | Metabolomics Work-bench | | |
| ST000483 | metabo-lomics | 40 | 40 | Metabolomics Work-bench | | |
| ST000484 | metabo-lomics | 41 | 40 | Metabolomics Work-bench | | |
| ST000485 | metabo-lomics | 6 | 560 | Metabolomics Work-bench | | |
| ST000496 | metabo-lomics | 69 | 100 | Metabolomics Work-bench | | |
| ST000608 | metabo-lomics | 264 | 60 | Metabolomics Work-bench | | |
| ST000659 | metabo-lomics | 15 | 138 | Metabolomics Work-bench | | |
| ST000805 | metabo-lomics | 18 | 60 | Metabolomics Work-bench | | |
| ST000808 | metabo-lomics | 45 | 60 | Metabolomics Work-bench | | |
| ST000843 | metabo-lomics | 86 | 67 | Metabolomics Work-bench | | |
| ST000872 | metabo-lomics | 74 | 66 | Metabolomics Work-bench | | |
| ST000884 | metabo-lomics | 727 | 183 | Metabolomics Work-bench | | |
| GSE48472 | methyla-tion | 485512 | 56 | BioDataome | various tissues | |
| GSE60185 | methyla-tion | 485512 | 285 | BioDataome | breast cancer | disease of cellular prolif-eration |

| | | | | | | |
|---|---|---|---|---|---|---|
| GSE61441 | methyla-tion | 485512 | 92 | BioDataome | renal cell carcinoma | disease of cellular prolif-eration |
| GSE63409 | methyla-tion | 485512 | 74 | BioDataome | acute myeloid leukemia | disease of cellular prolif-eration |
| GSE63695 | methyla-tion | 485512 | 97 | BioDataome | osteoarthritis | disease of anatomical en-tity |
| GSE66210 | methyla-tion | 485512 | 60 | BioDataome | patau_edwards_and_down_syn-drome | genetic disease |
| GSE66313 | methyla-tion | 485512 | 55 | BioDataome | breast cancer | disease of cellular prolif-eration |
| GSE66552 | methyla-tion | 485512 | 45 | BioDataome | deletion or duplication of 7q11.23 | disease of mental health |
| GSE66836 | methyla-tion | 485512 | 183 | BioDataome | lung cancer | disease of cellular prolif-eration |
| GSE68060 | methyla-tion | 485512 | 120 | BioDataome | | disease of cellular prolif-eration |
| GSE68825 | methyla-tion | 485512 | 144 | BioDataome | lung cancer | disease of cellular prolif-eration |
| GSE68838 | methyla-tion | 485512 | 302 | BioDataome | colon cancer | disease of cellular prolif-eration |
| GSE69502 | methyla-tion | 485512 | 179 | BioDataome | neural tube defects | physical disorder |
| GSE71955 | methyla-tion | 485512 | 135 | BioDataome | graves' disease | disease of anatomical en-tity |
| GSE71957 | methyla-tion | 485512 | 135 | BioDataome | graves' disease | disease of anatomical en-tity |
| GSE72867 | methyla-tion | 485512 | 69 | BioDataome | viral infectious disease | |
| GSE72872 | methyla-tion | 485512 | 250 | BioDataome | barrett's esophagus | disease of anatomical en-tity |
| GSE72874 | methyla-tion | 485512 | 250 | BioDataome | barrett's esophagus | disease of anatomical en-tity |
| GSE74167 | methyla-tion | 485512 | 42 | BioDataome | obesity | disease of metabolism |

| | | | | | | |
|---|---|---|---|---|---|---|
| GSE74432 | methyla-tion | 485512 | 122 | BioDataome | sotos syndrome | genetic disease |
| GSE74738 | methyla-tion | 485512 | 79 | BioDataome | imprinted methylation in pla-centa | disease of anatomical en-tity |
| GSE77269 | methyla-tion | 485512 | 60 | BioDataome | hepatocellular carcinoma | disease of cellular prolif-eration |
| GSE77276 | methyla-tion | 485512 | 60 | BioDataome | hepatocellular carcinoma | disease of cellular prolif-eration |
| GSE77348 | methyla-tion | 485512 | 85 | BioDataome | breast cancer | disease of cellular prolif-eration |
| GSE77954 | methyla-tion | 485512 | 48 | BioDataome | colorectal adenocarcinoma | disease of cellular prolif-eration |
| GSE77955 | methyla-tion | 485512 | 48 | BioDataome | colorectal adenocarcinoma | disease of cellular prolif-eration |
| GSE79695 | methyla-tion | 485512 | 44 | BioDataome | acute myeloid leukemia | disease of cellular prolif-eration |
| GSE80468 | methyla-tion | 485512 | 60 | BioDataome | | disease of anatomical en-tity |
| GSE85210 | methyla-tion | 485512 | 253 | BioDataome | cigarette smoke exposure | |
| GSE85828 | methyla-tion | 485512 | 114 | BioDataome | induced pluripotency | |
| GSE10006 | microar-ray | 54675 | 87 | BioDataome | chronic obstructive pulmonary disease | disease of anatomical en-tity |
| GSE10041 | microar-ray | 54675 | 72 | BioDataome | relaxation response | |
| GSE10063 | microar-ray | 54675 | 60 | BioDataome | lung cancer | disease of cellular prolif-eration |
| GSE10135 | microar-ray | 54675 | 59 | BioDataome | cancer | disease of cellular prolif-eration |
| GSE10334 | microar-ray | 54675 | 247 | BioDataome | aggressive periodontitis | disease of anatomical en-tity |
| GSE10780 | microar-ray | 54675 | 185 | BioDataome | breast cancer | disease of cellular prolif-eration |

| | | | | | | |
|---|---|---|---|---|---|---|
| GSE10810 | microar-ray | 54675 | 58 | BioDataome | breast cancer | disease of cellular prolif-eration |
| GSE10890 | microar-ray | 54675 | 112 | BioDataome | breast carcinoma | disease of cellular prolif-eration |
| GSE10927 | microar-ray | 54675 | 65 | BioDataome | adrenocortical carcinoma | disease of cellular prolif-eration |
| GSE11083 | microar-ray | 54675 | 83 | BioDataome | atherosclerosis | disease of anatomical en-tity |
| GSE11135 | microar-ray | 54675 | 204 | BioDataome | leukemia | disease of cellular prolif-eration |
| GSE11190 | microar-ray | 54675 | 78 | BioDataome | hepatitis c | disease by infectious agent |
| GSE11348 | microar-ray | 54675 | 93 | BioDataome | viral infectious disease | disease by infectious agent |
| GSE11755 | microar-ray | 54675 | 41 | BioDataome | sepsis | disease by infectious agent |
| GSE11784 | microar-ray | 54675 | 171 | BioDataome | chronic obstructive pulmonary disease | disease of anatomical en-tity |
| GSE11869 | microar-ray | 54675 | 75 | BioDataome | endometrial cancer | disease of cellular prolif-eration |
| GSE11906 | microar-ray | 54675 | 217 | BioDataome | pulmonary emphysema | disease of anatomical en-tity |
| GSE11952 | microar-ray | 54675 | 83 | BioDataome | chronic obstructive pulmonary disease | disease of anatomical en-tity |
| GSE12452 | microar-ray | 54675 | 41 | BioDataome | nasopharyngeal carcinoma | disease of cellular prolif-eration |
| GSE12453 | microar-ray | 54675 | 67 | BioDataome | non-hodgkin lymphoma | disease of cellular prolif-eration |
| GSE12460 | microar-ray | 54675 | 64 | BioDataome | neuroblastoma | disease of cellular prolif-eration |
| GSE12662 | microar-ray | 54675 | 106 | BioDataome | acute promyelocytic leukemia | disease of cellular prolif-eration |
| GSE12711 | microar-ray | 54675 | 47 | BioDataome | | |

| | | | | | | |
|---|---|---|---|---|---|---|
| GSE12790 | microar-ray | 54675 | 98 | BioDataome | breast cancer | disease of cellular prolif-eration |
| GSE13139 | microar-ray | 54675 | 54 | BioDataome | atherosclerosis | disease of anatomical en-tity |
| GSE13355 | microar-ray | 54675 | 180 | BioDataome | psoriasis | disease of anatomical en-tity |
| GSE13367 | microar-ray | 54675 | 56 | BioDataome | ulcerative colitis | disease of anatomical en-tity |
| GSE13548 | microar-ray | 54675 | 42 | BioDataome | cancer | disease of cellular prolif-eration |
| GSE13732 | microar-ray | 54675 | 113 | BioDataome | multiple sclerosis | disease of anatomical en-tity |
| GSE13896 | microar-ray | 54675 | 70 | BioDataome | chronic obstructive pulmonary disease | disease of anatomical en-tity |
| GSE13911 | microar-ray | 54675 | 69 | BioDataome | gastric cancer | disease of cellular prolif-eration |
| GSE13931 | microar-ray | 54675 | 98 | BioDataome | chronic obstructive pulmonary disease | disease of anatomical en-tity |
| GSE13933 | microar-ray | 54675 | 87 | BioDataome | lung disease | disease of anatomical en-tity |
| GSE14671 | microar-ray | 54675 | 59 | BioDataome | chronic myeloid leukemia | disease of cellular prolif-eration |
| GSE14905 | microar-ray | 54675 | 82 | BioDataome | psoriasis | disease of anatomical en-tity |
| GSE14924 | microar-ray | 54675 | 41 | BioDataome | acute myeloid leukemia | disease of cellular prolif-eration |
| GSE15061 | microar-ray | 54675 | 870 | BioDataome | myelodysplastic syndrome | disease of cellular prolif-eration |
| GSE15396 | microar-ray | 54675 | 156 | BioDataome | | |
| GSE15490 | microar-ray | 54675 | 50 | BioDataome | chronic lymphocytic leukemia | disease of cellular prolif-eration |
| GSE15605 | microar-ray | 54675 | 74 | BioDataome | melanoma | disease of cellular prolif-eration |

| | | | | | | |
|---|---|---|---|---|---|---|
| GSE15645 | microar-ray | 54675 | 42 | BioDataome | arthropathy | disease of anatomical en-tity |
| GSE15913 | microar-ray | 54675 | 40 | BioDataome | chronic lymphocytic leukemia | disease of cellular prolif-eration |
| GSE16059 | microar-ray | 54675 | 88 | BioDataome | chronic fatigue syndrome | syndrome |
| GSE16134 | microar-ray | 54675 | 310 | BioDataome | periodontitis | disease of anatomical en-tity |
| GSE16214 | microar-ray | 54675 | 240 | BioDataome | multiple sclerosis | disease of anatomical en-tity |
| GSE16387 | microar-ray | 54675 | 46 | BioDataome | | |
| GSE16395 | microar-ray | 54675 | 48 | BioDataome | langerhans-cell histiocytosis | disease of anatomical en-tity |
| GSE16515 | microar-ray | 54675 | 52 | BioDataome | pancreatic cancer | disease of cellular prolif-eration |
| GSE16696 | microar-ray | 54675 | 45 | BioDataome | lung disease | disease of anatomical en-tity |
| GSE16837 | microar-ray | 54675 | 113 | BioDataome | bacterial infection | |
| GSE16879 | microar-ray | 54675 | 133 | BioDataome | crohn's disease | disease of anatomical en-tity |
| GSE17612 | microar-ray | 54675 | 51 | BioDataome | schizophrenia | disease of mental health |
| GSE17624 | microar-ray | 54675 | 60 | BioDataome | | disease of cellular prolif-eration |
| GSE17674 | microar-ray | 54675 | 62 | BioDataome | neuroectodermal tumor | disease of cellular prolif-eration |
| GSE17679 | microar-ray | 54675 | 117 | BioDataome | neuroectodermal tumor | disease of cellular prolif-eration |
| GSE17905 | microar-ray | 54675 | 157 | BioDataome | chronic obstructive pulmonary disease | disease of anatomical en-tity |
| GSE18105 | microar-ray | 54675 | 111 | BioDataome | colorectal cancer | disease of cellular prolif-eration |

| | | | | | | |
|---|---|---|---|---|---|---|
| GSE18123 | microar-ray | 54675 | 99 | BioDataome | autism spectrum disorder | disease of anatomical en-tity |
| GSE18206 | microar-ray | 54675 | 48 | BioDataome | irritant dermatitis | disease of anatomical en-tity |
| GSE18520 | microar-ray | 54675 | 63 | BioDataome | spindle cell carcinoma | disease of cellular prolif-eration |
| GSE18521 | microar-ray | 54675 | 75 | BioDataome | spindle cell carcinoma | disease of cellular prolif-eration |
| GSE18781 | microar-ray | 54675 | 55 | BioDataome | | disease of anatomical en-tity |
| GSE18791 | microar-ray | 54675 | 57 | BioDataome | viral infectious disease | disease by infectious agent |
| GSE18842 | microar-ray | 54675 | 91 | BioDataome | lung cancer | disease of cellular prolif-eration |
| GSE19069 | microar-ray | 54675 | 147 | BioDataome | adult T-cell leukemia | disease of cellular prolif-eration |
| GSE19123 | microar-ray | 54675 | 44 | BioDataome | breast cancer | disease of cellular prolif-eration |
| GSE19188 | microar-ray | 54675 | 156 | BioDataome | lung cancer | disease of cellular prolif-eration |
| GSE19407 | microar-ray | 54675 | 127 | BioDataome | chronic obstructive pulmonary disease | disease of anatomical en-tity |
| GSE19667 | microar-ray | 54675 | 121 | BioDataome | lung disease | disease of anatomical en-tity |
| GSE19722 | microar-ray | 54675 | 58 | BioDataome | lung adenocarcinoma | disease of cellular prolif-eration |
| GSE19743 | microar-ray | 54675 | 177 | BioDataome | | |
| GSE19804 | microar-ray | 54675 | 120 | BioDataome | non-small cell lung carcinoma | disease of cellular prolif-eration |
| GSE20257 | microar-ray | 54675 | 135 | BioDataome | chronic obstructive pulmonary disease | disease of anatomical en-tity |
| GSE20489 | microar-ray | 54675 | 54 | BioDataome | | |

| | | | | | | |
|---|---|---|---|---|---|---|
| GSE20713 | microar-ray | 54675 | 108 | BioDataome | breast cancer | disease of cellular prolif-eration |
| GSE20916 | microar-ray | 54675 | 145 | BioDataome | colorectal cancer | disease of cellular prolif-eration |
| GSE21094 | microar-ray | 54675 | 49 | BioDataome | acute lymphoblastic leukemia | genetic disease |
| GSE21138 | microar-ray | 54675 | 59 | BioDataome | schizophrenia | disease of mental health |
| GSE2125 | microar-ray | 54675 | 45 | BioDataome | b-cell lymphoma | disease of cellular prolif-eration |
| GSE21510 | microar-ray | 54675 | 104 | BioDataome | colorectal cancer | disease of cellular prolif-eration |
| GSE21545 | microar-ray | 54675 | 223 | BioDataome | atherosclerosis | disease of anatomical en-tity |
| GSE21912 | microar-ray | 54675 | 42 | BioDataome | multiple myeloma | disease of cellular prolif-eration |
| GSE22047 | microar-ray | 54675 | 230 | BioDataome | chronic obstructive pulmonary disease | disease of anatomical en-tity |
| GSE22229 | microar-ray | 54675 | 58 | BioDataome | kidney transplantation | |
| GSE22459 | microar-ray | 54675 | 65 | BioDataome | kidney transplantation | |
| GSE23293 | microar-ray | 54675 | 41 | BioDataome | chronic lymphocytic leukemia | disease of cellular prolif-eration |
| GSE23394 | microar-ray | 54675 | 96 | BioDataome | non-hodgkin lymphoma | disease of cellular prolif-eration |
| GSE23878 | microar-ray | 54675 | 59 | BioDataome | colorectal cancer | disease of cellular prolif-eration |
| GSE24147 | microar-ray | 54675 | 42 | BioDataome | leukemia | disease of cellular prolif-eration |
| GSE24223 | microar-ray | 54675 | 179 | BioDataome | | |
| GSE25902 | microar-ray | 54675 | 120 | BioDataome | | |

| | | | | | | |
|---|---|---|---|---|---|---|
| GSE26049 | microar-ray | 54675 | 181 | BioDataome | essential thrombocythemia | disease of cellular prolif-eration |
| GSE26378 | microar-ray | 54675 | 103 | BioDataome | toxic shock syndrome | disease by infectious agent |
| GSE26440 | microar-ray | 54675 | 130 | BioDataome | toxic shock syndrome | disease by infectious agent |
| GSE27202 | microar-ray | 54675 | 48 | BioDataome | | |
| GSE27210 | microar-ray | 54675 | 114 | BioDataome | | |
| GSE27536 | microar-ray | 54675 | 54 | BioDataome | | disease of anatomical en-tity |
| GSE27562 | microar-ray | 54675 | 162 | BioDataome | breast cancer | disease of cellular prolif-eration |
| GSE27567 | microar-ray | 54675 | 162 | BioDataome | breast cancer | disease of cellular prolif-eration |
| GSE27716 | microar-ray | 54675 | 40 | BioDataome | lung adenocarcinoma | disease of cellular prolif-eration |
| GSE27719 | microar-ray | 54675 | 40 | BioDataome | lung adenocarcinoma | disease of cellular prolif-eration |
| GSE27830 | microar-ray | 54675 | 155 | BioDataome | breast cancer | disease of cellular prolif-eration |
| GSE27854 | microar-ray | 54675 | 115 | BioDataome | colorectal cancer | disease of cellular prolif-eration |
| GSE27858 | microar-ray | 54675 | 56 | BioDataome | chronic lymphocytic leukemia | disease of cellular prolif-eration |
| GSE27913 | microar-ray | 54675 | 115 | BioDataome | colorectal cancer | disease of cellular prolif-eration |
| GSE28492 | microar-ray | 54675 | 80 | BioDataome | | |
| GSE28702 | microar-ray | 54675 | 54 | BioDataome | colorectal cancer | disease of cellular prolif-eration |
| GSE28750 | microar-ray | 54675 | 41 | BioDataome | sepsis | |

| | | | | | | |
|---|---|---|---|---|---|---|
| GSE29265 | microar-ray | 54675 | 49 | BioDataome | thyroid cancer | disease of cellular prolif-eration |
| GSE29431 | microar-ray | 54675 | 66 | BioDataome | breast cancer | disease of cellular prolif-eration |
| GSE30219 | microar-ray | 54675 | 307 | BioDataome | lung cancer | disease of cellular prolif-eration |
| GSE30240 | microar-ray | 54675 | 75 | BioDataome | cancer | disease of cellular prolif-eration |
| GSE30674 | microar-ray | 54675 | 56 | BioDataome | common cold | disease of anatomical en-tity |
| GSE30678 | microar-ray | 54675 | 56 | BioDataome | common cold | disease of anatomical en-tity |
| GSE30784 | microar-ray | 54675 | 229 | BioDataome | squamous cell carcinoma | disease of cellular prolif-eration |
| GSE31048 | microar-ray | 54675 | 221 | BioDataome | chronic lymphocytic leukemia | disease of cellular prolif-eration |
| GSE31189 | microar-ray | 54675 | 92 | BioDataome | bladder cancer | disease of anatomical en-tity |
| GSE31210 | microar-ray | 54675 | 246 | BioDataome | lung cancer | disease of cellular prolif-eration |
| GSE32448 | microar-ray | 54675 | 80 | BioDataome | prostate cancer | disease of cellular prolif-eration |
| GSE32474 | microar-ray | 54675 | 174 | BioDataome | cancer | disease of cellular prolif-eration |
| GSE33356 | microar-ray | 54675 | 120 | BioDataome | lung adenocarcinoma | disease of cellular prolif-eration |
| GSE33630 | microar-ray | 54675 | 105 | BioDataome | thyroid cancer | disease of cellular prolif-eration |
| GSE33943 | microar-ray | 54675 | 58 | BioDataome | inflammatory bowel disease | disease of anatomical en-tity |
| GSE34437 | microar-ray | 54675 | 66 | BioDataome | | |
| GSE34450 | microar-ray | 54675 | 132 | BioDataome | chronic obstructive pulmonary disease | disease of anatomical en-tity |

| | | | | | | |
|---|---|---|---|---|---|---|
| GSE34748 | microar-ray | 54675 | 56 | BioDataome | kidney transplantation | |
| GSE35570 | microar-ray | 54675 | 116 | BioDataome | thyroid cancer | disease of cellular prolif-eration |
| GSE35711 | microar-ray | 54675 | 49 | BioDataome | bacterial pneumonia | disease of anatomical en-tity |
| GSE35713 | microar-ray | 54675 | 203 | BioDataome | bacterial pneumonia | disease of anatomical en-tity |
| GSE35725 | microar-ray | 54675 | 114 | BioDataome | bacterial pneumonia | disease of anatomical en-tity |
| GSE35864 | microar-ray | 54675 | 72 | BioDataome | acquired immunodeficiency syn-drome | disease by infectious agent |
| GSE36769 | microar-ray | 54675 | 60 | BioDataome | breast cancer | disease of cellular prolif-eration |
| GSE36895 | microar-ray | 54675 | 76 | BioDataome | renal cell carcinoma | disease of cellular prolif-eration |
| GSE36907 | microar-ray | 54675 | 46 | BioDataome | chronic lymphocytic leukemia | disease of cellular prolif-eration |
| GSE37168 | microar-ray | 54675 | 40 | BioDataome | chronic lymphocytic leukemia | disease of cellular prolif-eration |
| GSE37364 | microar-ray | 54675 | 94 | BioDataome | colorectal cancer | disease of cellular prolif-eration |
| GSE37416 | microar-ray | 54675 | 48 | BioDataome | bacterial infection | |
| GSE37964 | microar-ray | 54675 | 44 | BioDataome | colorectal cancer | disease of cellular prolif-eration |
| GSE38663 | microar-ray | 54675 | 52 | BioDataome | hepatitis c | disease by infectious agent |
| GSE38666 | microar-ray | 54675 | 45 | BioDataome | ovarian cancer | disease of cellular prolif-eration |
| GSE39088 | microar-ray | 54675 | 142 | BioDataome | systemic lupus erythematosus | disease of anatomical en-tity |
| GSE39156 | microar-ray | 54675 | 64 | BioDataome | thyroid cancer | disease of cellular prolif-eration |

| | | | | | | |
|---|---|---|---|---|---|---|
| GSE39411 | microar-ray | 54675 | 152 | BioDataome | chronic lymphocytic leukemia | disease of cellular prolif-eration |
| GSE39612 | microar-ray | 54675 | 138 | BioDataome | merkel cell carcinoma | disease of cellular prolif-eration |
| GSE40611 | microar-ray | 54675 | 49 | BioDataome | Sjogren's syndrome | disease of anatomical en-tity |
| GSE40791 | microar-ray | 54675 | 194 | BioDataome | lung cancer | disease of cellular prolif-eration |
| GSE41296 | microar-ray | 54675 | 120 | BioDataome | cancer | disease of cellular prolif-eration |
| GSE41662 | microar-ray | 54675 | 48 | BioDataome | psoriasis | disease of anatomical en-tity |
| GSE41663 | microar-ray | 54675 | 81 | BioDataome | psoriasis | disease of anatomical en-tity |
| GSE41664 | microar-ray | 54675 | 157 | BioDataome | psoriasis | disease of anatomical en-tity |
| GSE41804 | microar-ray | 54675 | 40 | BioDataome | hepatocellular carcinoma | disease of cellular prolif-eration |
| GSE41828 | microar-ray | 54675 | 50 | BioDataome | spindle cell sarcoma | disease of cellular prolif-eration |
| GSE41831 | microar-ray | 54675 | 78 | BioDataome | arthropathy | disease of anatomical en-tity |
| GSE41861 | microar-ray | 54675 | 138 | BioDataome | asthma | disease of anatomical en-tity |
| GSE41862 | microar-ray | 54675 | 116 | BioDataome | asthma | disease of anatomical en-tity |
| GSE42045 | microar-ray | 54675 | 50 | BioDataome | spindle cell carcinoma | disease of cellular prolif-eration |
| GSE42048 | microar-ray | 54675 | 62 | BioDataome | nonpapillary renal cell carci-noma | disease of cellular prolif-eration |
| GSE42050 | microar-ray | 54675 | 194 | BioDataome | nonpapillary renal cell carci-noma | disease of cellular prolif-eration |
| GSE42057 | microar-ray | 54675 | 136 | BioDataome | | disease of anatomical en-tity |

| | | | | | | |
|---|---|---|---|---|---|---|
| GSE42204 | microar-ray | 54675 | 65 | BioDataome | b-cell lymphoma | disease of cellular prolif-eration |
| GSE42568 | microar-ray | 54675 | 121 | BioDataome | breast cancer | disease of cellular prolif-eration |
| GSE42743 | microar-ray | 54675 | 103 | BioDataome | oral cavity cancer | disease of cellular prolif-eration |
| GSE4290 | microar-ray | 54675 | 180 | BioDataome | neuroectodermal tumor | disease of cellular prolif-eration |
| GSE43010 | microar-ray | 54675 | 42 | BioDataome | cancer | disease of cellular prolif-eration |
| GSE4302 | microar-ray | 54675 | 118 | BioDataome | asthma | disease of anatomical en-tity |
| GSE43081 | microar-ray | 54675 | 97 | BioDataome | melanoma | disease of cellular prolif-eration |
| GSE43346 | microar-ray | 54675 | 68 | BioDataome | lung cancer | disease of cellular prolif-eration |
| GSE43939 | microar-ray | 54675 | 149 | BioDataome | chronic obstructive pulmonary disease | disease of anatomical en-tity |
| GSE44927 | microar-ray | 54675 | 40 | BioDataome | prostate cancer | disease of cellular prolif-eration |
| GSE45267 | microar-ray | 54675 | 87 | BioDataome | spindle cell carcinoma | disease of cellular prolif-eration |
| GSE45404 | microar-ray | 54675 | 42 | BioDataome | colorectal cancer | disease of cellular prolif-eration |
| GSE45426 | microar-ray | 54675 | 45 | BioDataome | response to exercise | |
| GSE45436 | microar-ray | 54675 | 134 | BioDataome | spindle cell carcinoma | disease of cellular prolif-eration |
| GSE45536 | microar-ray | 54675 | 123 | BioDataome | rheumatoid arthritis | disease of anatomical en-tity |
| GSE45537 | microar-ray | 54675 | 135 | BioDataome | rheumatoid arthritis | disease of anatomical en-tity |
| GSE45827 | microar-ray | 54675 | 155 | BioDataome | cancer | disease of cellular prolif-eration |

| | | | | | | |
|---|---|---|---|---|---|---|
| GSE45867 | microar-ray | 54675 | 40 | BioDataome | rheumatoid arthritis | disease of anatomical en-tity |
| GSE4607 | microar-ray | 54675 | 123 | BioDataome | | |
| GSE46474 | microar-ray | 54675 | 40 | BioDataome | kidney transplantation | |
| GSE46699 | microar-ray | 54675 | 130 | BioDataome | renal cell carcinoma | disease of metabolism |
| GSE47908 | microar-ray | 54675 | 60 | BioDataome | ulcerative colitis | disease of anatomical en-tity |
| GSE48311 | microar-ray | 54675 | 56 | BioDataome | endotoxinic challenge | |
| GSE49695 | microar-ray | 54675 | 64 | BioDataome | chronic lymphocytic leukemia | disease of cellular prolif-eration |
| GSE49697 | microar-ray | 54675 | 64 | BioDataome | chronic lymphocytic leukemia | disease of cellular prolif-eration |
| GSE50006 | microar-ray | 54675 | 220 | BioDataome | chronic lymphocytic leukemia | disease of cellular prolif-eration |
| GSE50161 | microar-ray | 54675 | 130 | BioDataome | medulloblastoma | disease of cellular prolif-eration |
| GSE50772 | microar-ray | 54675 | 81 | BioDataome | systemic lupus erythematosus | disease of anatomical en-tity |
| GSE50811 | microar-ray | 54675 | 241 | BioDataome | endometrial cancer | disease of cellular prolif-eration |
| GSE50830 | microar-ray | 54675 | 168 | BioDataome | endometrial cancer | disease of cellular prolif-eration |
| GSE50831 | microar-ray | 54675 | 189 | BioDataome | endometrial cancer | disease of cellular prolif-eration |
| GSE51024 | microar-ray | 54675 | 96 | BioDataome | lung cancer | disease of cellular prolif-eration |
| GSE51401 | microar-ray | 54675 | 64 | BioDataome | hepatocellular carcinoma | disease of cellular prolif-eration |
| GSE51981 | microar-ray | 54675 | 148 | BioDataome | endometriosis | disease of anatomical en-tity |

| | | | | | | |
|---|---|---|---|---|---|---|
| GSE52237 | microar-ray | 54675 | 58 | BioDataome | lung disease | disease of anatomical entity |
| GSE52553 | microar-ray | 54675 | 84 | BioDataome | alcohol dependence | disease of mental health |
| GSE52724 | microar-ray | 54675 | 286 | BioDataome | diabetes insipidus | disease of anatomical entity |
| GSE5281 | microar-ray | 54675 | 161 | BioDataome | alzheimer's disease | disease of anatomical entity |
| GSE53091 | microar-ray | 54675 | 125 | BioDataome | colorectal cancer | disease of cellular proliferation |
| GSE5350 | microar-ray | 54675 | 140 | BioDataome | spindle cell carcinoma | disease of cellular proliferation |
| GSE53552 | microar-ray | 54675 | 99 | BioDataome | psoriasis | disease of anatomical entity |
| GSE53757 | microar-ray | 54675 | 144 | BioDataome | renal cell carcinoma | disease of cellular proliferation |
| GSE53987 | microar-ray | 54675 | 205 | BioDataome | schizophrenia | disease of mental health |
| GSE54129 | microar-ray | 54675 | 132 | BioDataome | | disease of cellular proliferation |
| GSE54169 | microar-ray | 54675 | 75 | BioDataome | mantle cell lymphoma | disease of cellular proliferation |
| GSE54219 | microar-ray | 54675 | 155 | BioDataome | breast cancer | disease of cellular proliferation |
| GSE55092 | microar-ray | 54675 | 140 | BioDataome | virus associated cancer | disease by infectious agent |
| GSE55201 | microar-ray | 54675 | 81 | BioDataome | psoriasis | disease of anatomical entity |
| GSE55849 | microar-ray | 54675 | 64 | BioDataome | chronic granulomatous disease | disease of anatomical entity |
| GSE56315 | microar-ray | 54675 | 122 | BioDataome | b-cell lymphoma | disease of cellular proliferation |
| GSE57051 | microar-ray | 54675 | 40 | BioDataome | arsenic exposure | |

| | | | | | | |
|---|---|---|---|---|---|---|
| GSE5816 | microar-ray | 54675 | 42 | BioDataome | colon cancer | disease of cellular prolif-eration |
| GSE58294 | microar-ray | 54675 | 92 | BioDataome | cardioembolic stroke | disease of anatomical en-tity |
| GSE58331 | microar-ray | 54675 | 175 | BioDataome | orbital inflammatory diseases | disease of anatomical en-tity |
| GSE5900 | microar-ray | 54675 | 78 | BioDataome | multiple myeloma | disease of cellular prolif-eration |
| GSE59294 | microar-ray | 54675 | 40 | BioDataome | atopic dermatitis | disease of anatomical en-tity |
| GSE59312 | microar-ray | 54675 | 79 | BioDataome | hepatitis c | |
| GSE60542 | microar-ray | 54675 | 92 | BioDataome | thyroid cancer | disease of anatomical en-tity |
| GSE61629 | microar-ray | 54675 | 54 | BioDataome | cancer | disease of cellular prolif-eration |
| GSE62232 | microar-ray | 54675 | 91 | BioDataome | hepatocellular carcinoma | disease by infectious agent |
| GSE6338 | microar-ray | 54675 | 60 | BioDataome | anaplastic large cell lymphoma | disease of cellular prolif-eration |
| GSE63514 | microar-ray | 54675 | 128 | BioDataome | cervical cancer | disease of cellular prolif-eration |
| GSE64256 | microar-ray | 54675 | 125 | BioDataome | colorectal cancer | disease of cellular prolif-eration |
| GSE64258 | microar-ray | 54675 | 125 | BioDataome | colorectal cancer | disease of cellular prolif-eration |
| GSE64300 | microar-ray | 54675 | 42 | BioDataome | | |
| GSE64738 | microar-ray | 54675 | 45 | BioDataome | mithramycin treatment | disease of cellular prolif-eration |
| GSE64857 | microar-ray | 54675 | 81 | BioDataome | colorectal cancer | disease of cellular prolif-eration |
| GSE64913 | microar-ray | 54675 | 59 | BioDataome | asthma | disease of anatomical en-tity |

| | | | | | | |
|---|---|---|---|---|---|---|
| GSE64951 | microar-ray | 54675 | 94 | BioDataome | gastric cancer | disease of cellular prolif-eration |
| GSE65010 | microar-ray | 54675 | 48 | BioDataome | rheumatoid arthritis | disease of anatomical en-tity |
| GSE65127 | microar-ray | 54675 | 40 | BioDataome | vitiligo | disease of anatomical en-tity |
| GSE65194 | microar-ray | 54675 | 178 | BioDataome | breast cancer | disease of cellular prolif-eration |
| GSE65216 | microar-ray | 54675 | 178 | BioDataome | breast cancer | disease of cellular prolif-eration |
| GSE65914 | microar-ray | 54675 | 58 | BioDataome | rosacea | disease of anatomical en-tity |
| GSE66272 | microar-ray | 54675 | 54 | BioDataome | cancer | disease of cellular prolif-eration |
| GSE66354 | microar-ray | 54675 | 149 | BioDataome | neuroectodermal tumor | disease of cellular prolif-eration |
| GSE66360 | microar-ray | 54675 | 99 | BioDataome | acute myocardial infarction | disease of anatomical en-tity |
| GSE67596 | microar-ray | 54675 | 72 | BioDataome | arthropathy | disease of anatomical en-tity |
| GSE6764 | microar-ray | 54675 | 75 | BioDataome | cancer | disease of cellular prolif-eration |
| GSE6791 | microar-ray | 54675 | 84 | BioDataome | virus associated cancer | disease of cellular prolif-eration |
| GSE68015 | microar-ray | 54675 | 112 | BioDataome | cancer | disease of cellular prolif-eration |
| GSE68801 | microar-ray | 54675 | 122 | BioDataome | alopecia areata | disease of anatomical en-tity |
| GSE71065 | microar-ray | 54675 | 80 | BioDataome | acquired immunodeficiency syn-drome | disease by infectious agent |
| GSE71222 | microar-ray | 54675 | 152 | BioDataome | colorectal cancer | disease of cellular prolif-eration |
| GSE71717 | microar-ray | 54675 | 60 | BioDataome | ishikawa cells treated with genistein | disease of cellular prolif-eration |

| | | | | | | |
|---|---|---|---|---|---|---|
| GSE71730 | microar-ray | 54675 | 47 | BioDataome | ulcerative colitis | disease of anatomical en-tity |
| GSE71996 | microar-ray | 54675 | 65 | BioDataome | | |
| GSE72140 | microar-ray | 54675 | 48 | BioDataome | melasma | disease of anatomical en-tity |
| GSE72754 | microar-ray | 54675 | 52 | BioDataome | systemic lupus erythematosus | disease of anatomical en-tity |
| GSE72798 | microar-ray | 54675 | 82 | BioDataome | lupus nephritis | disease of anatomical en-tity |
| GSE72925 | microar-ray | 54675 | 168 | BioDataome | kidney disease | disease of anatomical en-tity |
| GSE75132 | microar-ray | 54675 | 41 | BioDataome | cervical cancer | disease of cellular prolif-eration |
| GSE7753 | microar-ray | 54675 | 47 | BioDataome | arthropathy | disease of anatomical en-tity |
| GSE77658 | microar-ray | 54675 | 184 | BioDataome | | |
| GSE77659 | microar-ray | 54675 | 184 | BioDataome | | |
| GSE7904 | microar-ray | 54675 | 62 | BioDataome | breast carcinoma | disease of cellular prolif-eration |
| GSE80060 | microar-ray | 54675 | 206 | BioDataome | | disease of anatomical en-tity |
| GSE8157 | microar-ray | 54675 | 43 | BioDataome | polycystic ovary syndrome | syndrome |
| GSE83556 | microar-ray | 54675 | 40 | BioDataome | fmr1 expansion | |
| GSE8507 | microar-ray | 54675 | 141 | BioDataome | job's syndrome | disease of anatomical en-tity |
| GSE8545 | microar-ray | 54675 | 54 | BioDataome | chronic obstructive pulmonary disease | disease of anatomical en-tity |
| GSE8581 | microar-ray | 54675 | 58 | BioDataome | cleft lip | physical disorder |

| | | | | | | |
|---|---|---|---|---|---|---|
| GSE86574 | microar-ray | 54675 | 110 | BioDataome | neuroectodermal tumor | disease of cellular prolif-eration |
| GSE8671 | microar-ray | 54675 | 64 | BioDataome | colorectal cancer | disease of cellular prolif-eration |
| GSE9196 | microar-ray | 54675 | 53 | BioDataome | | |
| GSE9348 | microar-ray | 54675 | 82 | BioDataome | colorectal cancer | disease of cellular prolif-eration |
| GSE94349 | microar-ray | 54675 | 168 | BioDataome | spinal meningioma | disease of cellular prolif-eration |
| GSE94521 | microar-ray | 54675 | 60 | BioDataome | developmental toxicity | disease of anatomical en-tity |
| GSE9489 | microar-ray | 54675 | 60 | BioDataome | kidney disease | disease of anatomical en-tity |
| GSE9493 | microar-ray | 54675 | 82 | BioDataome | kidney disease | disease of anatomical en-tity |
| GSE9692 | microar-ray | 54675 | 45 | BioDataome | toxic shock syndrome | disease by infectious agent |
| GSE9826 | microar-ray | 54675 | 45 | BioDataome | ovarian cancer | disease of cellular prolif-eration |
| SRP007825 | rnaseq | 50794 | 66 | BioDataome | psoriasis | disease of anatomical en-tity |
| SRP010038 | rnaseq | 38320 | 40 | BioDataome | pterygium | disease of anatomical en-tity |
| SRP022043 | rnaseq | 21030 | 70 | BioDataome | alzheimer's disease | disease of anatomical en-tity |
| SRP023262 | rnaseq | 54197 | 100 | BioDataome | breast cancer | disease of cellular prolif-eration |
| SRP024274 | rnaseq | 44498 | 160 | BioDataome | chronic obstructive pulmonary disease | disease of anatomical en-tity |
| SRP026042 | rnaseq | 51433 | 84 | BioDataome | psoriasis | disease of anatomical en-tity |
| SRP029880 | rnaseq | 50053 | 54 | BioDataome | colorectal cancer | disease of cellular prolif-eration |

| | | | | | | |
|---|---|---|---|---|---|---|
| SRP035988 | rnaseq | 55315 | 188 | BioDataome | psoriasis | disease of anatomical entity |
| SRP041036 | rnaseq | 49766 | 60 | BioDataome | multiple myeloma | disease of cellular proliferation |
| SRP041538 | rnaseq | 54954 | 187 | BioDataome | chronic obstructive pulmonary disease | disease of anatomical entity |
| SRP043162 | rnaseq | 43274 | 53 | BioDataome | asthma | disease of anatomical entity |
| SRP044668 | rnaseq | 53677 | 99 | BioDataome | glioma | disease of cellular proliferation |
| SRP047194 | rnaseq | 53473 | 54 | BioDataome | autism spectrum disorder | disease of mental health |
| SRP049097 | rnaseq | 57743 | 54 | BioDataome | leiomyosarcoma | disease of cellular proliferation |
| SRP049988 | rnaseq | 51327 | 102 | BioDataome | lung disease | |
| SRP050331 | rnaseq | 41297 | 192 | BioDataome | prostate cancer | disease of cellular proliferation |
| SRP051368 | rnaseq | 45070 | 185 | BioDataome | bacterial infection | |
| SRP051848 | rnaseq | 51338 | 187 | BioDataome | post-traumatic stress disorder | disease of mental health |
| SRP053794 | rnaseq | 50311 | 40 | BioDataome | atopic dermatitis | disease of anatomical entity |
| SRP056220 | rnaseq | 38214 | 309 | BioDataome | trophoblast differentiation | |
| SRP056733 | rnaseq | 49093 | 152 | BioDataome | bacterial infection | disease by infectious agent |
| SRP059057 | rnaseq | 47438 | 73 | BioDataome | coeliac disease | disease of anatomical entity |
| SRP061881 | rnaseq | 52834 | 224 | BioDataome | juvenile idiopathic arthritis | |
| SRP062025 | rnaseq | 53915 | 73 | BioDataome | myelodysplastic syndrome | disease of cellular proliferation |
| SRP062966 | rnaseq | 46792 | 117 | BioDataome | systemic lupus erythematosus | disease of anatomical entity |
| SRP063867 | rnaseq | 51094 | 95 | BioDataome | induced pluripotency | disease of anatomical entity |

| | | | | | | |
|---|---|---|---|---|---|---|
| SRP065812 | rnaseq | 56965 | 52 | BioDataome | psoriasis | disease of anatomical entity |
